# KIF23 regulation by miR-107 controls replicative tumor cell fitness in mouse and human hepatocellular carcinoma

**DOI:** 10.1101/2023.11.20.565448

**Authors:** Mirco Castoldi, Sanchari Roy, Carolin Lohr, Rossella Pellegrino, Mihael Vucur, Michael T. Singer, Veronika Buettner, Matthias A. Dille, Lara R. Heij, Lars Zender, Ulf P. Neumann, Thomas Longerich, Christoph Roderburg, Tom Luedde

## Abstract

**Background & Aims:** In hepatocellular carcinoma, there is a lack of successful translation of experimental targets identified in mouse models to human patients. In this study, we used a comprehensive transcriptomic approach in mice to identify novel potential targets for therapeutic intervention in humans.

**Methods:** We analysed combined genome-wide miRNA and mRNA expression data in three pathogenically distinct mouse models of liver cancer. Effects of target genes on hepatoma cell fitness were evaluated by proliferation, survival and motility assays. TCGA and GEO databases, in combination with tissue microarrays (TMA), were used to validate the mouse targets and their impact on human HCC prognosis. Finally, the functional effects of the identified targets on tumorigenesis and tumor therapy were tested in hydrodynamic tail vein injection (HDTVi)-based preclinical HCC models *in vivo*.

**Results:** The expression of miR-107 was found to be significantly reduced in mouse models of liver tumors of various etiologies and in cohorts of human HCC patients. Overexpression of miR-107 or inhibition of its novel target Kinesin family member 23 (Kif23) significantly reduced proliferation by interfering with cytokinesis, thereby controlling survival and motility of mouse and human hepatoma cells. In humans, KIF23 expression was found to be a prognostic marker in liver cancer, with high expression associated with poor prognosis. HDTVi of vectors carrying either pre-miR- 107 or anti-Kif23 shRNA inhibited the development of highly aggressive cMyc-NRas-induced liver cancers in mice.

**Conclusions:** Disruption of the miR-107/Kif23 axis inhibited hepatoma cell proliferation *in vitro* and prevented oncogene-induced liver cancer development *in vivo*, offering a novel potential avenue for the treatment of HCC in humans.

**Impact and implications:** A comprehensive analysis integrating *in silico* prediction, miRNA and mRNA data in three pathogenically distinct mouse models provided novel targets for the treatment of human HCC, bridging the translational gap between mouse data and human HCC. Our functional findings on the novel miR-107/Kif23 module provide important new insights into the control of mitosis in liver cancer cells. The findings that miR-107 overexpression or Kif23 inhibition had a dramatic functional effect on inhibiting the growth of liver cancer cells *in vitro* and *in vivo* suggest that the miR-107/Kif23 axis may be a promising novel target and potential adjunct to sequential systemic therapy of HCC.

**Graphical abstract:** 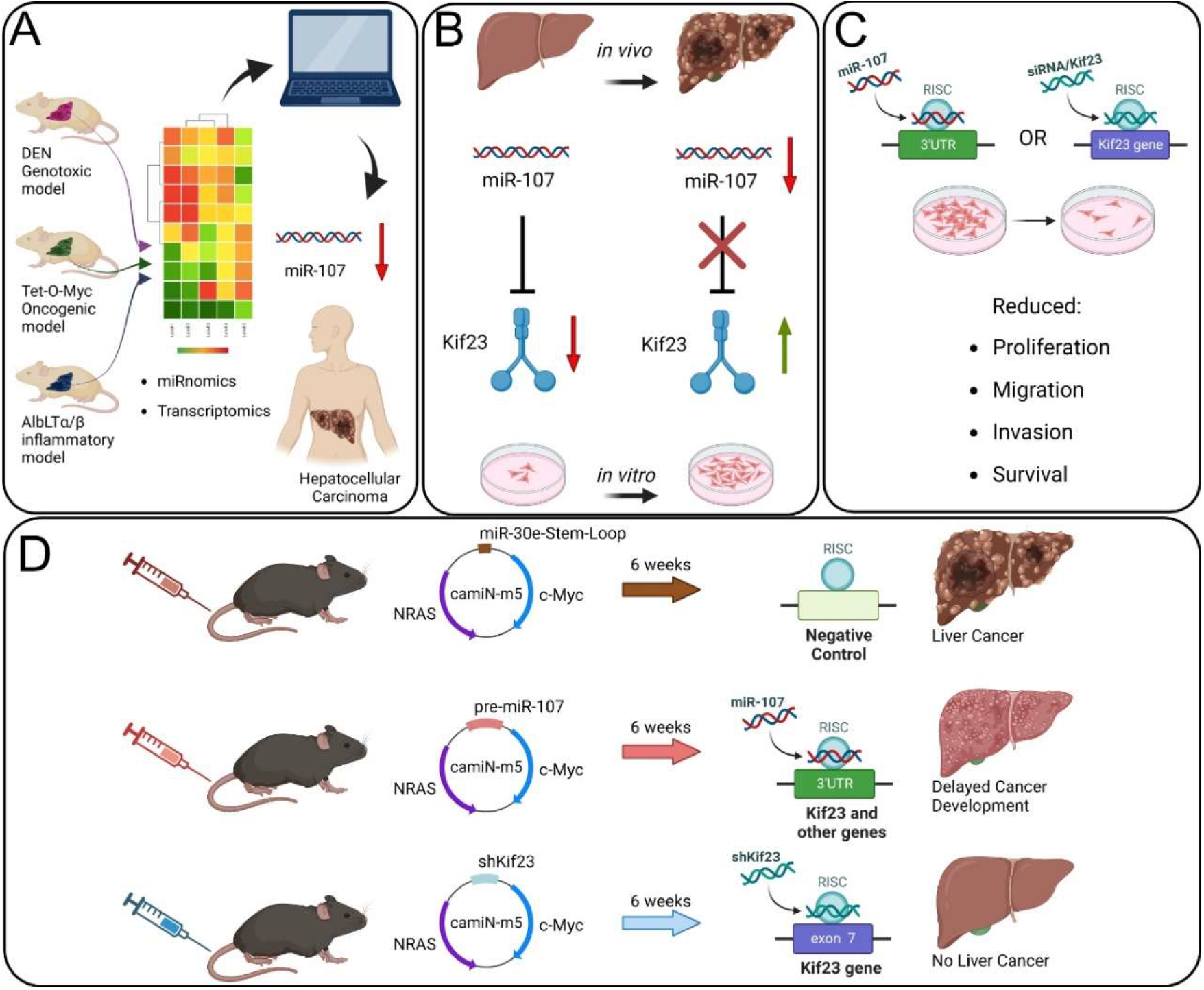

**Highlights:**

- miR-107 is globally downregulated in mouse liver cancers of different etiologies and represents a potential biomarker in human HCC.
- Integration of *in-silico*-prediction, miRNA and mRNA transcriptomics identified KIF23, a mitotic spindle-associated protein, as a specific target mediating the biological effects of miR-107.
- The miR-107/KIF23 module promotes replicative fitness of liver cancer cells through an essential function in cytokinesis
- Mice receiving shRNA targeting Kif23 were completely protected from oncogene-induced liver cancer.

## Introduction

Hepatocellular carcinoma (HCC) is the most common type of primary liver cancer [1]. Despite advancements in identifying risk factors, comprehending disease etiology, and developing anti-viral strategies against viral hepatitis, the incidence of HCC is on the rise. Therapeutic options are limited and survival after diagnosis is poor, in fact, the incidence of HCC still nearly equals its mortality rate [2]. Therefore, to effectively combat HCC, a more comprehensive genetic and mechanistic understanding is crucial, highlighting the need for further research and improved preventive measures.

MicroRNAs (miRNAs) are key epigenetic regulators of the transcriptome, whose dysregulation has been linked to increased fitness of cancer cells and to the control of oncogenic hallmarks [3]. Although a large and growing body of literature indicates that dysregulation of miRNA expression plays an important role in hepatocarcinogenesis [4], the rate of successful translation of miRNA-centered animal studies to clinical trials has been hampered by an inadequate ability to predict the human response to treatment. HCC in humans is characterized by wide inter- and intra-tumor heterogeneity, and because individual animal models can replicate only some molecular aspects of human HCC pathogenesis, it is currently unclear which animal model of liver cancer is best suited for studying molecular mechanisms of significant relevance to humans. In addition, it must be considered that miRNAs regulate cellular physiology by fine tuning gene expression within larger regulatory networks [5]. Thus, although miRNAs themselves are not the functional mediators, their analysis may reveal relevant genes that could serve as potential therapeutic targets.

To identify dysregulated miRNA networks relevant to human liver cancer development, systems biology approaches were used to analyze and correlate the miRnome and transcriptome in mouse models of liver cancer [6]. In the present study, we combined *in silico* prediction with comprehensive miRNA and mRNA transcriptomic analysis and identified miR-107 as globally downregulated in HCC. We found that miR-107 controls the migration and invasion of liver cancer cells through the fine-tuning of KIF23, a spindle-associated motor protein critical for cytokinesis [7], representing a promising biomarker and therapeutic target in human HCC.

## Materials and Methods

### Data retrieval from the Genome Omnibus repository and analysis

The miRNA and mRNA profiling data in animal model of liver cancer have been retrieved from GEO [8] under the super-series accession number GSE102418. This super-series is composed of the following sub-series: GSE102416, which contains the raw data from the analysis of miRNAs expression, and GSE102417 that contains the CEL files from the Affymetrix microarray. CEL files were analyzed by using the comprehensive transcriptome analysis software AltAnalyze [9], with fold changes cutoff ≥ 2; p ≤ 0.05. For the purpose of this analysis, arrays from mice treated with DEN, AlbLTα/β and Tet-O-Myc were grouped as normal, non-tumor (NTL) and tumor liver, regardless of pathogenesis. Enrichment analysis was performed with ShinyGO [10] by uploading list of significantly upregulated genes, which were predicted to be miR-107 targets by miRWalk [11] target prediction database. Intersect between the microarray data and the miRWalk-generated list of miR-107 predicted targets was identified by using Venny Venn diagram (https://bioinfogp.cnb.csic.es/tools/venny/).

### Tissue Microarrays

A tissue microarray (TMA) containing non-tumorous and tumor liver tissues from HCC patients (n= 62; Patient’s characteristics in **Supplementary Table 1**) was constructed as previously described [12]. Tissue samples were provided by the Institute of Pathology, University Hospital Aachen and their use was approved by the local ethics committee in compliance with the Helsinki Declaration (vote number EK122/16). According to the vote, informed consent was not required because only long-term archived (>5 years) anonymized samples were used. Immunohistochemistry was performed on 3-µm sections. Antigen retrieval was carried out using citrate buffer solution pH 6 (Dako, Glostrup, Denmark). Detection was performed using the EnVision method (Dako) and counterstaining was done with hemalum solution. Staining was assessed using the immunoreactive score as previously described [13]: Primary antibody used to stain the TMAs: anti-KIF23 (1:100, Abcam Cambridge, MA, USA, ab174304). TMA scoring: 0, absent; 1–4, weak; 5–8, moderate; 9–12, strong expression.

### Analyses of the Cancer Genome Atlas (TCGA) Data and Kaplan-Meier Survival Curves

Some of the result shown in this study are based upon data generated by the TCGA Research Network: https://www.cancer.gov/tcga (accessed on 7th of April 2023). Kaplan–Meier survival curves of LIHC patients for miR-107, Kif23, PRC1, NUF2, AURKB, ANLN, and TTK were generated by using the KM Plotter [14].

### Statistical Analysis and Imaging Software

Statistical analyses were carried out using GraphPad Prism (version 9.4.1). To evaluate whether samples were normally distributed, the D’Agostino and Pearson normality tests were carried out. When the sample distribution passed the normality test, then parametric tests were carried out (i.e., one-way analysis of variance/ANOVA for three or more samples and a two-tailed t-test for two samples). When the samples did not pass the normality test, non-parametric tests were applied (i.e., the Kruskal– Wallis test for three or more samples and the Mann–Whitney test for two samples). The data were considered significant at a p value ≤ 0.05. Images were prepared using Affinity Designer (version 1.10.4.1198). Data were analyzed using GraphPad Prism (GraphPad Software, Inc., La Jolla, CA, Version 9.4.1).

### Ethics approval

The relevant official authority for animal protection (Landesamt für Natur, Umwelt und Verbraucherschutz Nordrhein-Westfalen, Recklinghausen, Germany) approved this study (reference number 84-02.04.2011.A133). Animals received care according to the German animal welfare act. Animal experiments included in this study were carried out in compliance with the Act No 246/1992 Coll., on the protection of animals against cruelty.

The ethics committee of the Medical Faculty of RWTH Aachen gave the approval (ethical votes EK122/16 and EK 206/09) for using human HCC tissue samples, in conformity with the Declaration of Helsinki ethical guidelines.

### Informed Consent Statement

All LIHC cohort patients who were involved in the TME preparation gave informed consent.

### Data Availability Statement

All data used in this study have been included in the manuscript.

## Results

### miR-107 is globally downregulated in mouse and human liver cancer

We analyzed genome-wide data analysis of miRNA expression in liver cancers of mice from a previously performed experiment including three prototypical models for liver cancer formation [i.e., Diethylnitrosamine (DEN) as a model of genotoxic hepatocarcinogenesis; Transgenic over-expression of Lymphotoxin (AlbLTα/β) as an inflammatory HCC model; and the oncogene-driven c-Myc (Tet-O-Myc mice); [6]]. This analysis revealed a common downregulation of miR-107 as a miRNA with a presently unclear function in hepatocarcinogenesis (**Figure 1A**). First, we measured and validated the downregulation of miR-107 expression in the livers of the three mouse models used in the genome-wide analysis (**Figure 1B and 1C**). We also corroborated these results by measuring the expression of miR-107 in the tumor tissue of other well-established mouse genetic models of liver cancer (namely: Tak1^LPC-KO^ [15, 16], Mcl-1^Δhep^ [17], Mdr2^KO^ [18] and Traf2/Ripk1^LPC-KO^ [19] mice; **Figure 1D**), and found that miR-107 was significantly downregulated in the tumors of all these respective animal models of HCC that we had analyzed. Finally, we assessed miR-107 levels in various mouse (Hepa 1-6, Hepa1c1c7) and human (Huh-7, HepG2) hepatoma cell lines and compared them to the expression in normal mouse and human livers, respectively.

**Figure 1.**
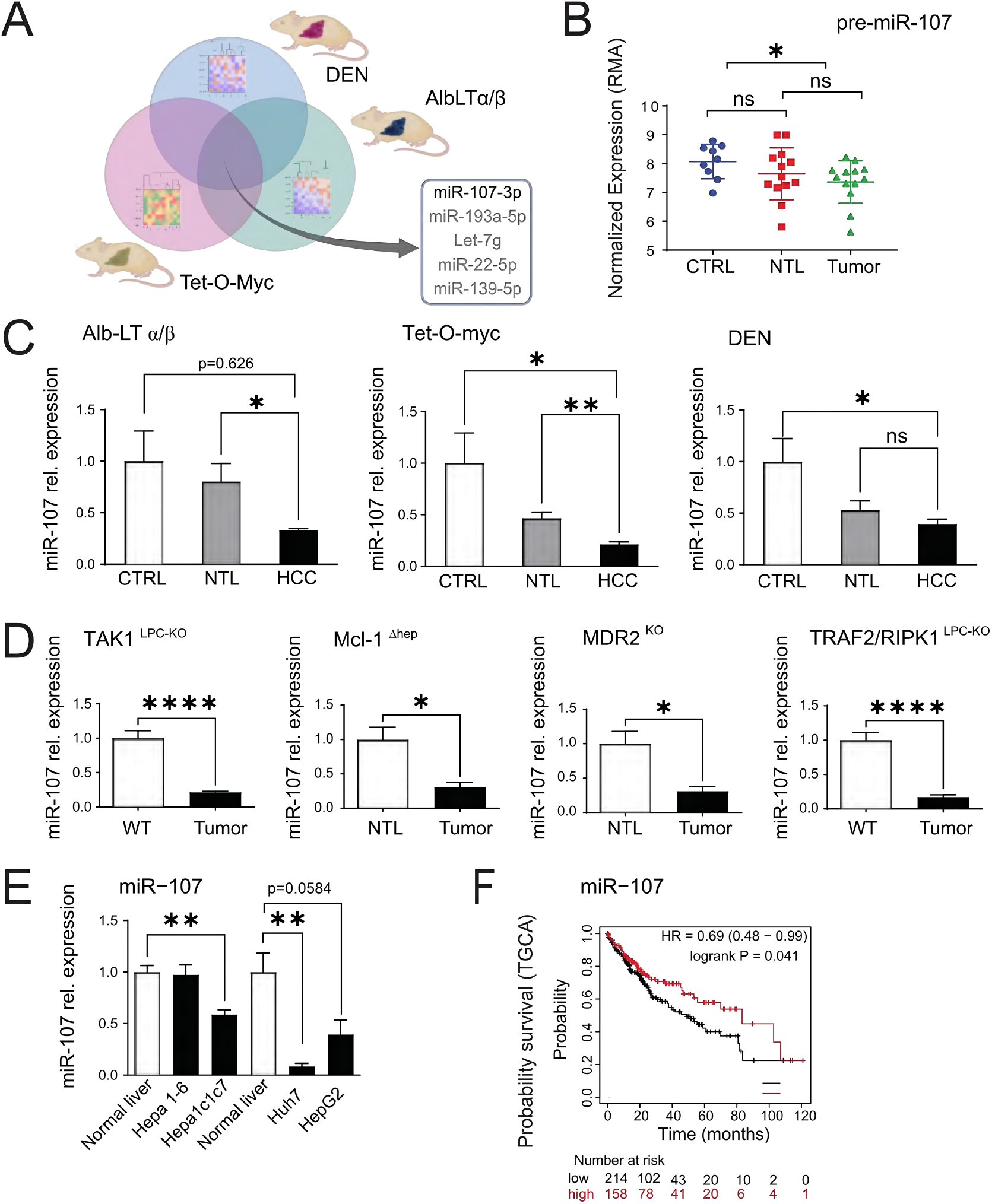
miR-107 is globally downregulated in etiologically unrelated mouse model of liver cancer and in human. (**A**) Systems biology was employed to globally identify miRNAs that were downregulated in the tumor tissues of three etiologically distinct mouse models of liver cancer: DEN, AlbLTα/β, and Tet-O-Myc. (**B**) RMA- normalized microarray data for the expression of pre-miR-107 in the healthy (n = 9), nontumoral liver (NTL: n = 13) and tumor (n = 13) liver tissue from the mice included in this study. (**C**) RT-qPCR validation of miR-107 expression in the individual DEN (Untreated, n = 6; NTL, n = 5; Tumor, n = 5), AlbLTα/β (Untreated, n = 4; NTL, n = 4; Tumor, n = 4), and Tet-O-Myc (Untreated, n = 4; NTL, n = 4; Tumor, n = 4) models of liver cancer. (**D**) RT-qPCR analysis of miR-107 in four different genetic HCC mouse tumor models: Mcl-1^Δhep^ (n = 5), TAK1-KO (n = 11), TRAF2-RIPK1-KO (n = 7), Mdr2- KO (n = 7) compared to either NTL or Wild type (WT) mice. (**E**) RT-qPCR of miR-107 expression in normal mouse (n = 5) and human (n = 3) livers and in mouse (Hepa1-6; n = 5, Hepa1c1c7; n = 5) and human hepatoma cells (Huh-7; n = 5, HepG2; n = 5, Hep3B; n = 5). (**F**) Kaplan-Meier curves for the overall survival of HCC patients plotted against time (months) based on miR-107 expression levels (TCGA LIHC cohort datasets; n = 372). Results are represented as mean ± SD, significant differences were evaluated by using 1-way ANOVA with Newman-Keuls post-hoc test or 2- tailed, unpaired t test t-test as appropriate. (*, p ≤ 0.05; **, p ≤ 0.001; ****, p ≤ 0.0001).

Consistent with the findings in murine liver cancer models, we observed a significant downregulation of miR-107 in both mouse Hepa1c1c7 and human Huh-7 hepatoma cell lines when compared to their respective controls (**Figure 1E**). To further explore the possible correlation between liver cancer development and miR-107 expression in humans, we conducted an analysis using liver hepatocellular carcinoma cohorts (LIHC) sourced from “The Cancer Genome Atlas” (TCGA) database. By employing Kaplan-Meier curves to assess patient survival, our study revealed a significant association linking decreased expression of miR-107 to significantly shorter survival in LIHC patients (**Figure 1F**). These findings underscore the potential significance of miR-107 in the progression of hepatocellular carcinoma in human.

### miR-107 upregulation negatively impacts the replicative fitness of human and mouse hepatoma cells

The discovery that miR-107 is downregulated in human and mouse models of liver cancer suggests either an “active” role in the development of liver cancer or a “passive” modulation resulting from underlying diseases. To unveil the role of miR-107 in liver cancer, we assessed its impact on the oncogenic hallmarks of cancer cells, such as cellular proliferation, survival, and motility. The activity of miR-107 was evaluated in human Huh-7 hepatoma cells and we found that miR-107 overexpression significantly inhibited the cells’ ability to proliferate and migrate, ultimately reducing their overall survival (**Figure 2A - 2G**). Specifically, we found that increased levels of miR-107 impaired Huh-7 colony formation (**Figure 2A**), proliferation (**Figures 2B and C**), and motility (**Figures 2E, F, and G**), correlating with increased cell death (**Figure 2D**). Overall, these results suggest that high levels of miR-107 significantly reduce the overall fitness of human hepatoma cells.

**Figure 2.**
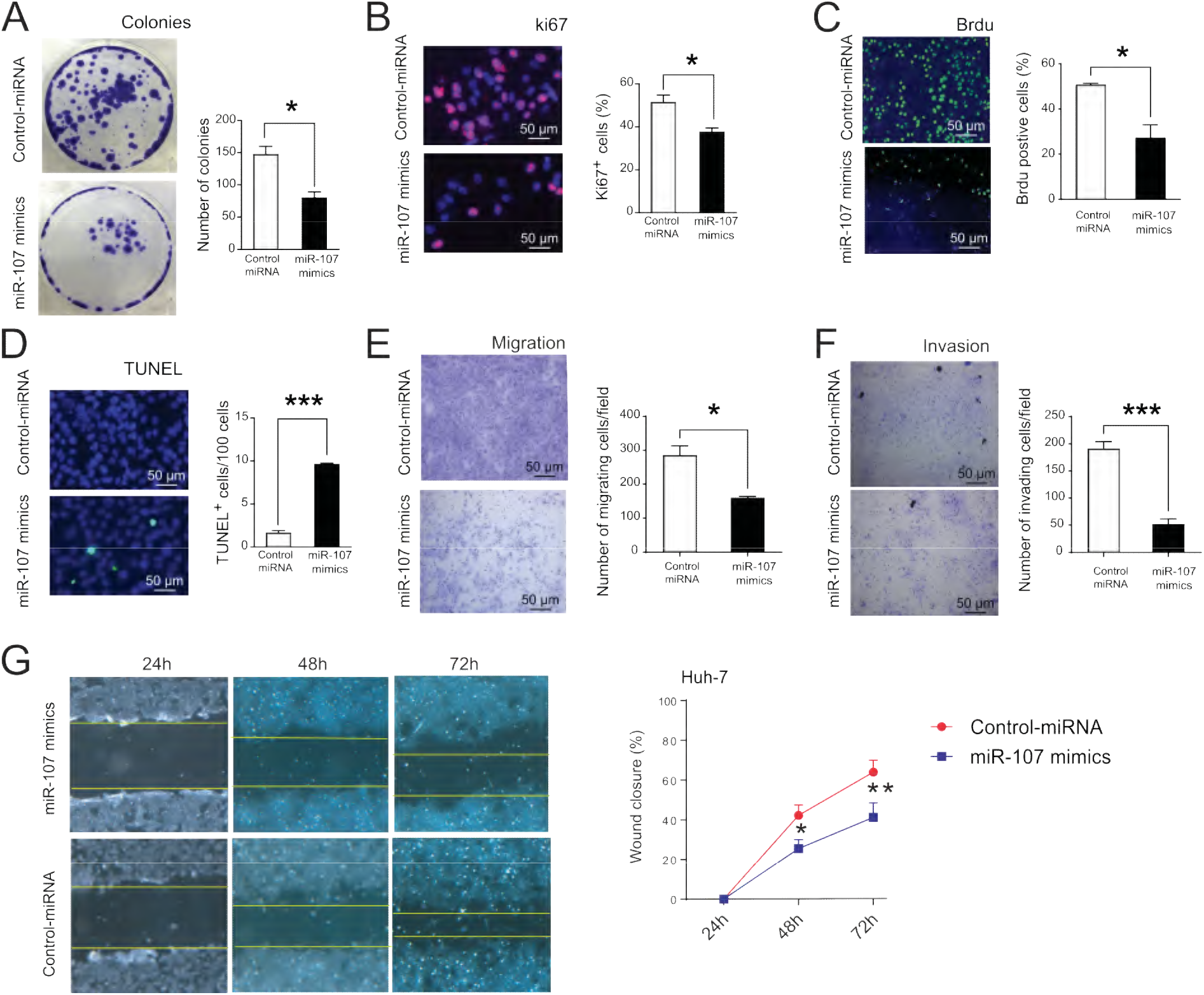
miR-107 upregulation, negatively impact the fitness of human hepatoma cells. Huh7 cells were transfected with either miR-107 mimic or control miRNA and the effect of miR-107 upregulation on cancer hallmark of hepatoma cells was assessed by (**A**) colony formation assay (n = 4), (**B**) Ki67 staining (n = 5), (**C**) BrdU proliferation assay (n = 5), (**D**) while Huh-7 viability was measured by TUNEL assay (n = 5). Transwell migration assay E) or Transwell coated with Matrigel invasion assay (**F**) for Hepa1c1c7 cells was determined after transfection with miR-107 mimic or control miRNAs for 72 h (n = 3). (**G**) Wound-healing assay was performed on Huh7 cells transfected with miR-107 mimic and wound closure was monitored over the indicated time post-transfection after treatment with Mitomycin C for 2 h (n = 3). Representative images are shown for the different experiments. Results are represented as mean ± SD, significant differences were evaluated by using 1-way ANOVA with Newman-Keuls post-hoc test or 2- tailed, unpaired t test t-test as appropriate. (*, p ≤ 0.05; **, p ≤ 0.01; ***, p ≤ 0.001).

The function of miR-107 was also evaluated in Hepa1c1c7 murine hepatoma cells. Overexpression of miR-107 in murine cells yielded outcomes comparable to those observed in human cells, including similar effects on cellular proliferation, survival, and motility (**Supplementary** Figure 1). These data support the conclusion that, in addition to inhibiting cell proliferation and inducing cell death, miR-107 overexpression hampers the migration and invasion capabilities of hepatoma cells.

### *In silico* prediction and integration with transcriptome data identified the mitotic spindle associated protein KIF23 as downstream target of miR-107

To identify miR-107-responsive genes whose upregulation may contribute to liver cancer development, we analyzed transcriptomic data from the livers of the DEN, AlbLTα/β, and Tet-O-Myc models (GSE102417 [6]; as outlined in the supplementary material). This led to the identification of 60 genes significantly upregulated in tumor tissue compared to both control and NTL tissues (**Figure 3A and Supplementary Table 2**). We then cross-referenced these genes with the predicted miR-107 targets from the miRWalk target prediction algorithm [11]), generating a list of 23 putative miR- 107 targets (**Supplementary Table 3**). Next, we loaded this gene list into gene-set enrichment tools [i.e., g:Profiler [20] and ShinyGo [10]) and performed gene set enrichment analysis (GSEA). The GSEA results showed that the miR-107 putative target genes had good enrichment score and significance in the cell cycle and mitotic spindle pathways (**Figure 3B and Supplementary** Figure 2A). Furthermore, GSEA identified that several of these genes may be an indicator of prognostic factors in the development of hepatocellular carcinoma (**Supplementary** Figure 2B).

**Figure 3.**
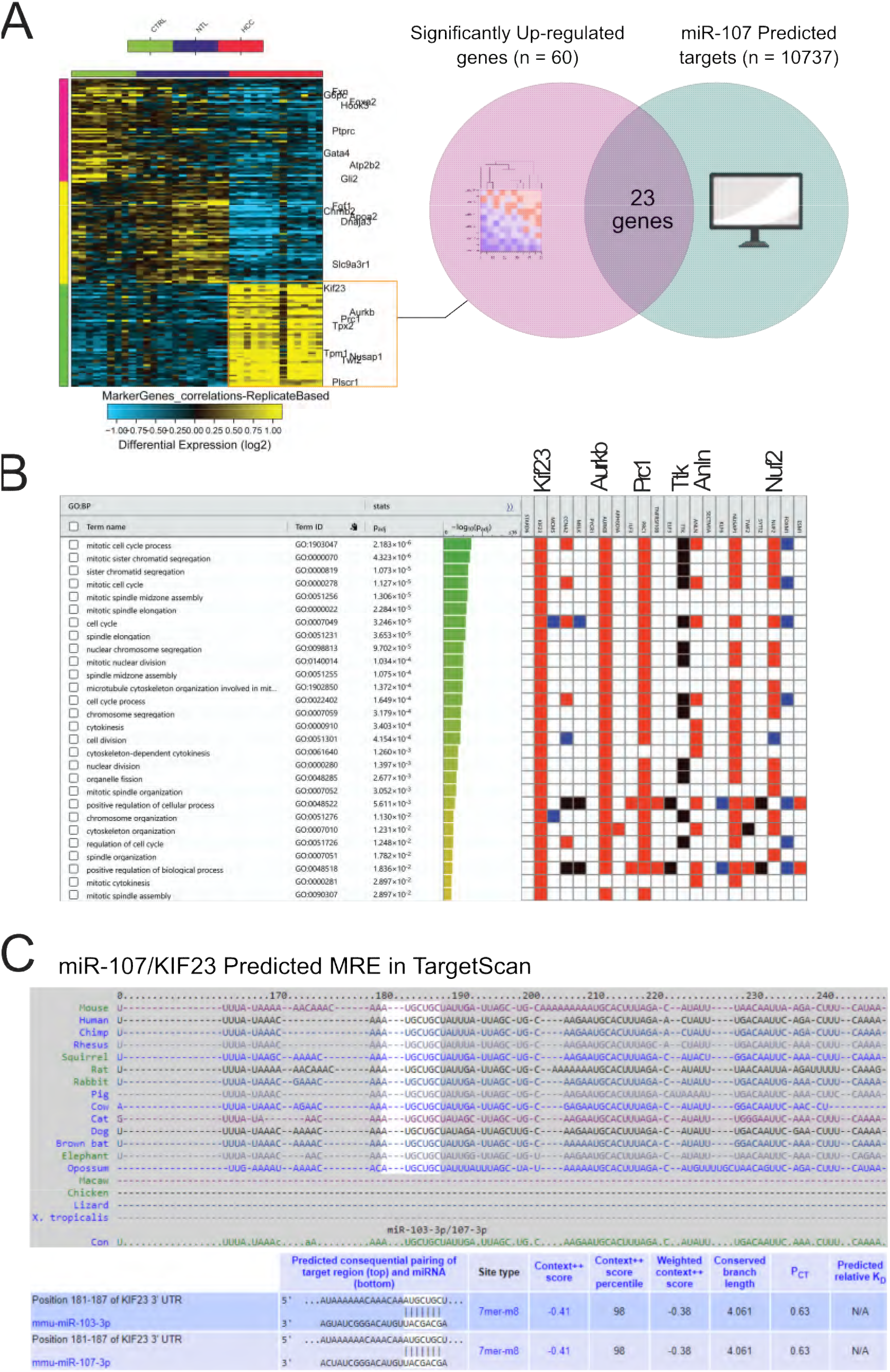
Kif23, a kinesin involved in cytokinesis, is a conserved direct target of miR-107. (A) Unsupervised hierarchical clustering of the genes regulated in the tumor tissue of the three mouse models used in this study (i.e., DEN, AlbLTα/β and Tet-O- Myc) compared to NTL and healthy tissue. The color scale illustrates the fold change of mRNAs across the samples (Generated by Altanalyze). The overlap between significantly upregulated genes and miRWalk predicted miR-107 targets is shown. (**B**) G:Profiler enrichment analysis of potential miR-107 targets and the name of the most significantly enriched genes (i.e. with lowest p-value) is given (**C**) TargetScan predicted the presence of a highly conserved binding site for miR-107 in the 3UTRs of KIF23 across different organism, unclosing human, mouse and rat. Results are represented as mean ± SD, significant differences were evaluated by 2- tailed, unpaired t test t-test. (*, p ≤ 0.05; ***, p ≤ 0.001; ****, p ≤ 0.0001).

To focus on the most promising candidates, the six most significantly enriched genes in the GSEA analysis were selected for further analysis: Kif23 (Kinesin family member 23), Aurkb (Aurora Kinase B), Prc1 (Polycomb repressive complex 1), Ttk (Monopolar spindle 1), Anln (Anillin), and Nuf2 (Nuclear Filament-containing protein 2). Our data suggest that these genes may play a role in liver cancer, given their significant upregulation in murine tumor tissue and their predicted involvement in mitotic spindle function.

To assess their relevance in humans, we queried the TCGA-LIHC cohort and performed individual Kaplan–Meier analyses to predict overall patient survival for each gene (**Supplementary** Figure 2C). Kaplan–Meier analyses of the TCGA-LIHC cohorts highlighted a negative correlation between the expression level of six genes and patient survival, suggesting Kif23, Aurkb, Prc1, Ttk, Anln, and Nuf2 have prognostic value in humans. Additionally, analysis of independent cohorts obtained from GEO (GSE6764 [21]; **Supplementary** Figure 2D) showed the significant upregulation of these genes in the livers of patients with advanced and very advanced HCC. Overall, these findings suggest that analyzing these six genes could provide more information about the molecular mechanism linking miR-107 dysregulation to liver cancer development and could potentially identify novel therapeutic targets for the treatment of HCC in humans.

### KIF23, a spindle-associated kinesin involved in cytokinesis, has a highly conserved miR-107 MRE in its 3’UTR

To focus our research on the most promising candidates, we used the TargetScan algorithm [22] to perform a conservation analysis of miR-107 responsive elements (MREs) in the 3’UTRs of Kif23, Aurkb, Prc1, Ttk, Anln, and Nuf2. TargetScan identifies putative MREs by searching for sequence conservation across multiple species in the region where the miRNA is predicted to bind. A high degree of sequence conservation suggests that the MRE is likely to be functional and that the gene is a potential target of the selected miRNA. Remarkably, among the six candidates only Kif23 exhibited a conserved MRE for miR-107 across multiple species (**Figure 3C**). KIF23, a spindle-associated kinesin crucial for cytokinesis [7], is found upregulated in several human malignancies, including colorectal cancer [23], pancreatic [24], ovarian cancers [25], and HCC [26]. We measured Kif23 levels in the three mouse models of HCC used in the transcriptomic analysis (i.e., DEN, AlbLTα/β, and Tet-O-Myc HCC models), and confirmed that Kif23 was significantly upregulated in the tumor tissues, but not in the non tumoral liver tissue (NTL) (**Supplementary** Figure 3A and B). To investigate the broader relevance of Kif23, we measured its expression in well-established murine liver cancer models (namely: Tak1^LPC-KO^ [15], Ikk2/Casp8^KO^ [27], Nemo/Ripk3^KO^ [27], Traf2/Casp8^LPC-KO^ [28] and Traf2/Ripk1^LPC-KO^ [19]), and found it to be significantly upregulated (**Supplementary** Figure 3C). To further explore the role of KIF23 in humans, we analyzed KIF23 expression in different human hepatoma cell lines at mRNA level (i.e., Huh-7, HepG2, and Hep3B) and found it to be significantly upregulated (**Supplementary** Figure 3D), confirming that KIF23 expression was dysregulated in both mice and human liver cancer cells.

### KIF23, is a direct target of miR-107, and promotes replicative fitness of liver cancer cells

To examine the subcellular localization of KIF23 in hepatoma cells, we performed immunofluorescence analysis with an anti-KIF23 antibody on Huh-7 cells (**Figure 4A**). We found that KIF23 is localized to the nucleus in interphase, relocates to the area of the microtubule organizing center of the mitotic spindle in dividing cells, and is mainly present in the spindle midzone during anaphase, where it is required for the assembly of the myosin contractile ring. To assess KIF23’s role in cytokinesis, we silenced KIF23 expression in Huh-7 cells using RNA interference (RNAi) and observed abnormal cell division, characterized by the formation of multinucleated cells (**Figure 4B**). These findings are consistent with a previous report [29].

**Figure 4.**
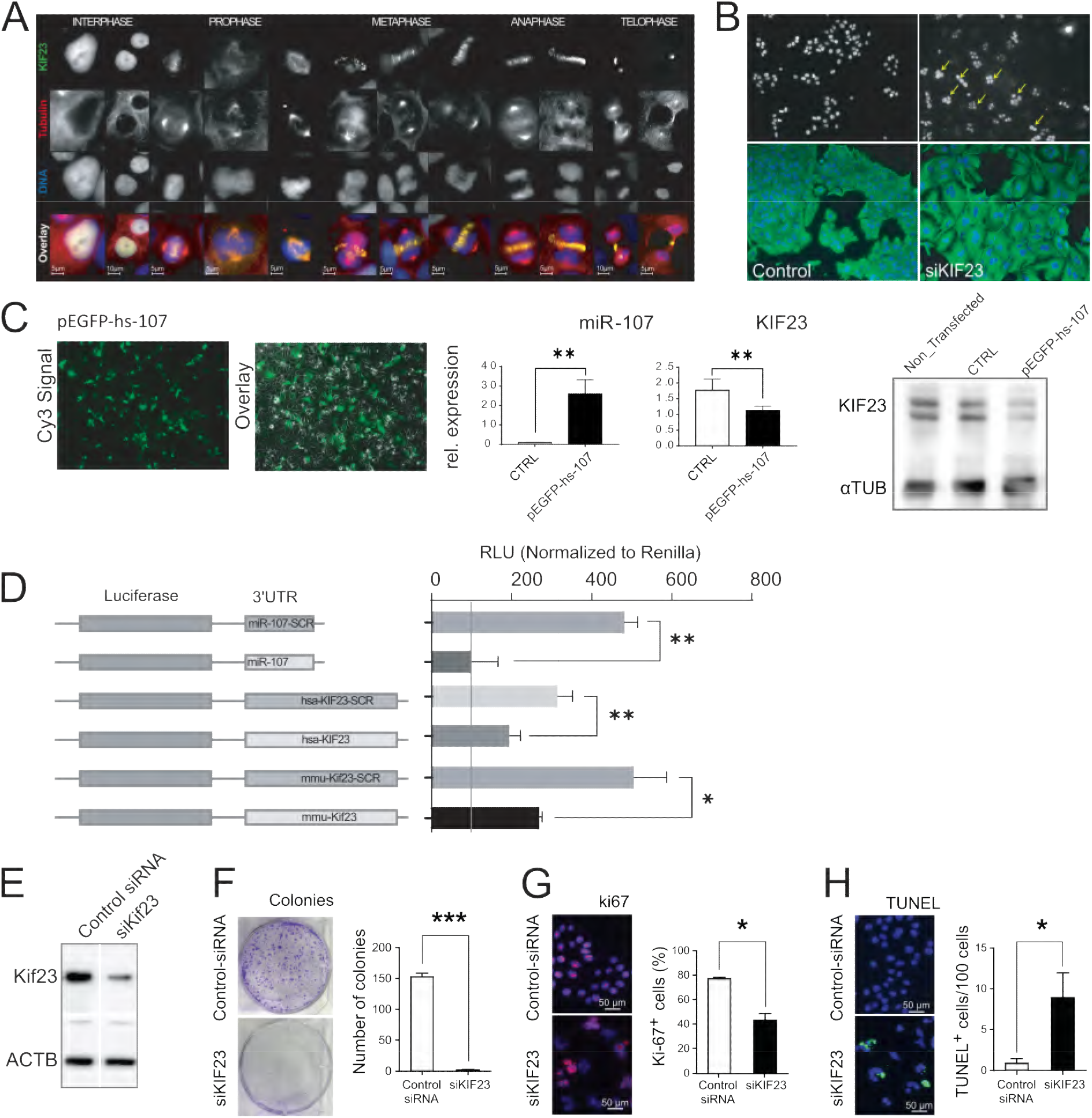
KIF23 silencing leads to incomplete cytokinesis and impairs survival of cancer cells. (**A**) Representative images showing KIF23 localization at the different stages of cell cycle (Interphase, Prophase, Metaphase, Anaphase and Telophase) in Huh-7 human hepatoma cells. (**B**) siRNA-mediated downregulation of KIF23 expression impairs cytokinesis, which results in cell division defects and in the formation of multinucleated cells. (**C**) KIF23 is significantly downregulated at mRNA and protein levels in Huh-7 cells transfected with an GFP vector encoding for miR107 (pEGFP-hs-107). (**D**) The full length 3UTRs for the mouse and human KIF23 were cloned into luciferase promoter vector in the sense or in the antisense orientation (SCR) as negative control. The plasmids miR107 containing the perfect miR-122 binding site in sense orientation, and miR107-SCR (miR-107 binding site in the antisense orientation) served as positive and negative control, respectively. Plasmids were transfected in presence or absence of miR-107 mimic into HEK293 cells and luciferase activity was measured 24 h post transfection. Relative luciferase activities are shown as percentage of the miR107 plasmid as transfection control (n = 4). (**E**) Huh-7 cells were transfected with either anti-Kif23 siRNAs or Control-siRNA and the effect inhibition on KIF23 was evaluated by Western Blot. Effect of KIF23 inhibition on Huh-7 fitness was assessed by (**F**) colony formation assay (n = 4), while Huh-7 viability and proliferation were respectively assessed by (**G**) TUNEL assay (n = 5) and (**H**) Ki67 staining (n = 5). Results are represented as mean ± SD, significant differences were evaluated by using 1-way ANOVA with Newman-Keuls post-hoc test or 2- tailed, unpaired t test as appropriate. (*, p ≤ 0.05; **, p ≤ 0.01; ***, p ≤ 0.001).

To investigate the potential interaction between miR-107 and KIF23, we transfected Huh-7 cells with a GFP vector encoding for human miR-107, resulting in a significant downregulation of KIF23 at both the RNA and protein levels (**Figure 4C**). These findings suggested that miR-107 regulates, directly or indirectly, the expression of KIF23. To determine the mechanism through which miR-107 regulates KIF23 expression, the full-length 3’UTRs of human and mouse KIF23 were cloned downstream of a luciferase reporter gene, either in the original (wild type, WT) or complementary reverse (CR) orientation (as described in [30]). Upon co-transfection of these constructs with miR-107 mimics into Huh-7 cells, a reduction in luciferase activity was observed specifically in the constructs carrying KIF23-3’UTRs in the native (i.e., WT) orientation. No such reduction was seen in the constructs carrying the 3’UTRs in the CR orientation (**Figure 4D**), unequivocally demonstrating that miR-107 directly regulates the expression of KIF23 in both human and mouse.

Next, the functional activity of KIF23 silencing on cellular proliferation and survival was investigated. Consistent with the phenotype observed in response to miR-107 overexpression, siRNA-mediated inhibition of KIF23 in Huh-7 cells (**Figure 4E**) led to a substantial decrease in cellular proliferation, as shown by colony formation (**Figure 4F**) and Ki67 assay (**Figure 4G**). Furthermore, KIF23 silencing led to a significant increase in the number of TUNEL-positive cells (**Figure 4H**). These findings support the conclusion that the dysregulation of the miR-107/KIF23 axis is involved in supporting HCC by enhancing tumor cell fitness in both mouse and human liver cancer.

### KIF23 nuclear localization is increased in human HCC tissue

Our data strongly support the conclusion that KIF23 is a pivotal functional mediator of miR-107, transmitting pro-carcinogenic signals in both murine and human liver cancers by supporting cancer cell fitness. While transcriptomic data provides valuable information about the cellular state and potential biomarkers, it is important to acknowledge that proteins are the primary agents responsible for cellular functions. Due to the influence of post-transcriptional machinery on translation and protein synthesis, discrepancies between measured mRNA levels and corresponding protein levels may arise. Therefore, to determine whether the increase in mRNA corresponded to an increase in protein level and to assess potential differences in KIF23 subcellular localization, KIF23 was analyzed in tissue microarrays (TMAs) containing cores from 62 different HCC patients [12]. TMAs were hybridized with an anti-KIF23 antibody, and the score associated with both KIF23 localization and signal intensity in the tumor tissue (HCC) and in the surrounding liver was calculated (**Figure 5, Supplementary Table 4;** [13]). In interphase KIF23 is expected to localize in the nucleus [31], however it has been shown that alternative splicing can generate a variant of KIF23 that is retained in the cytoplasm of cancer cells [32]. Along this line, KIF23 was detected in both cytosol and nuclei of liver cells, however, while no significant differences were observed in its cytoplasmic localization between tumor and non-tumor tissue, the assessment of KIF23’s nuclear localization identified a significant increase in KIF23 in the nuclei of the cells localized within the tumor compared to non-tumor tissues. These findings corroborate our analysis of publicly available datasets (**Supplementary** Figure 2) and suggest that increased nuclear localization of KIF23 may be a novel prognostic marker in human hepatocellular carcinoma.

**Figure 5.**
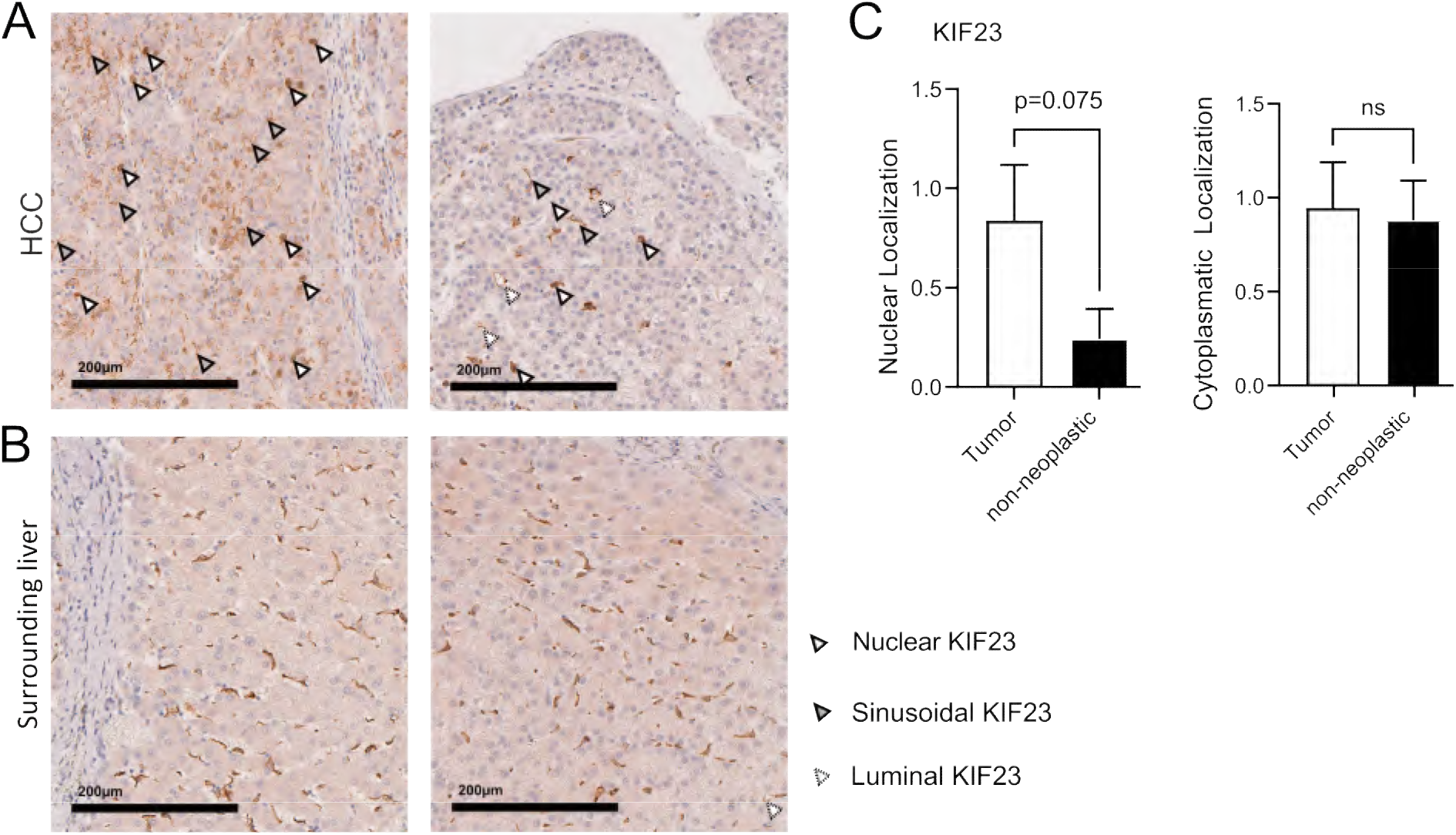
TMA analysis of KIF23 localization in the tumor compared to surrounding livers in HCC patients. Representative images of Tissue microarrays (TMA) prepared with cores of livers of human patients with HCC from (**A**) tumor tissue and (**B**) surrounding liver. KIF23 was detected in both cytosol and nuclei of liver cells. (**C**) Higher hepatocyte specific nuclear KIF23 expression (white arrow) was detected in the HCCs compared to the surrounding normal liver tissues, which instead showed mainly sinusoidal expression of KIF23 (grey arrow). Some HCCs displayed KIF23 luminal staining (dashed arrow), barely observed in surrounding liver. Weak expression of cytoplasmatic KIF23 was seen in both liver tissue types, without significant differences. Results are represented as mean ± SD, significant differences were evaluated by using 2- tailed, unpaired t test t-test as appropriate. (ns = Not significant).

### Targeting the miR-107/KIF23 axis inhibits oncogene-induced liver cancer formation

Given the pronounced impact of miR-107 and its target Kif23, on fundamental biological characteristics of liver cancer cells, our objective was to investigate whether manipulating their expressions could affect the progression and proliferation of liver cancer in a pre-clinical model. To induce liver cancer in mice, we employed hydrodynamic tail vein injection (HDTVi); to administer a transposon vector (camiN- mRE) encoding for the cMyc/NRas oncogenes which is known to produce highly aggressive hepatoma [33], in presence of either shRNA targeting KIF23, a pre-miR- 107, or a miR-30e backbone (negative control; miRE). The SplashRNA algorithm [34] was used to design three distinct shRNAs targeting various regions of murine Kif23 cDNA, which were then cloned into the RT3GEPIR entry vector [35]. Vectors carrying the shRNAs and the pre-miR-107 were transfected into murine Hepa1-6 hepatoma cells, and their efficacy in targeting Kif23 was assessed by Western Blot (**Supplementary** Figure 4). The most efficient shRNA in targeting Kif23 (shKif675), pri-miR107, and the miRE were cloned into the camiNmA vector as previously described [6]. The resulting constructs were co-injected with a vector expressing the transposase into the tail veins of 7-week-old male C57BL/6j mice using HDTVi. Animals were euthanized 6 weeks after injection, and their livers were examined for tumor presence.

Remarkably, no tumors were observed in animals injected with oncogenes along with either shKif675 or pre-miR-107 (**Figure 6A - D** and **Supplementary** Figure 5). In line with these findings, the liver-to-body-weight ratio (**Figure 6E** and **Supplementary Table 5**) and serum liver enzymes (i.e., **ALT,** and **AST; Figure 6F** and **Supplementary Table 6**) were only found to be significantly elevated in the miRE-injected, but not in the pre-miR-107 and shKif23-injected compared to untreated animals. Molecular analyses of selected markers (i.e., cMyc, NRAS, Kif23, and miR-107) performed by RT-qPCR from whole liver extracts showed a concordant regulation with tumor development in the respective mouse models (**Figure 6G**). Overall, the initial macroscopic analysis of the livers suggests that targeting the miR-107/Kif23 axis could be a promising new approach for the prevention or treatment of liver cancer.

**Figure 6.**
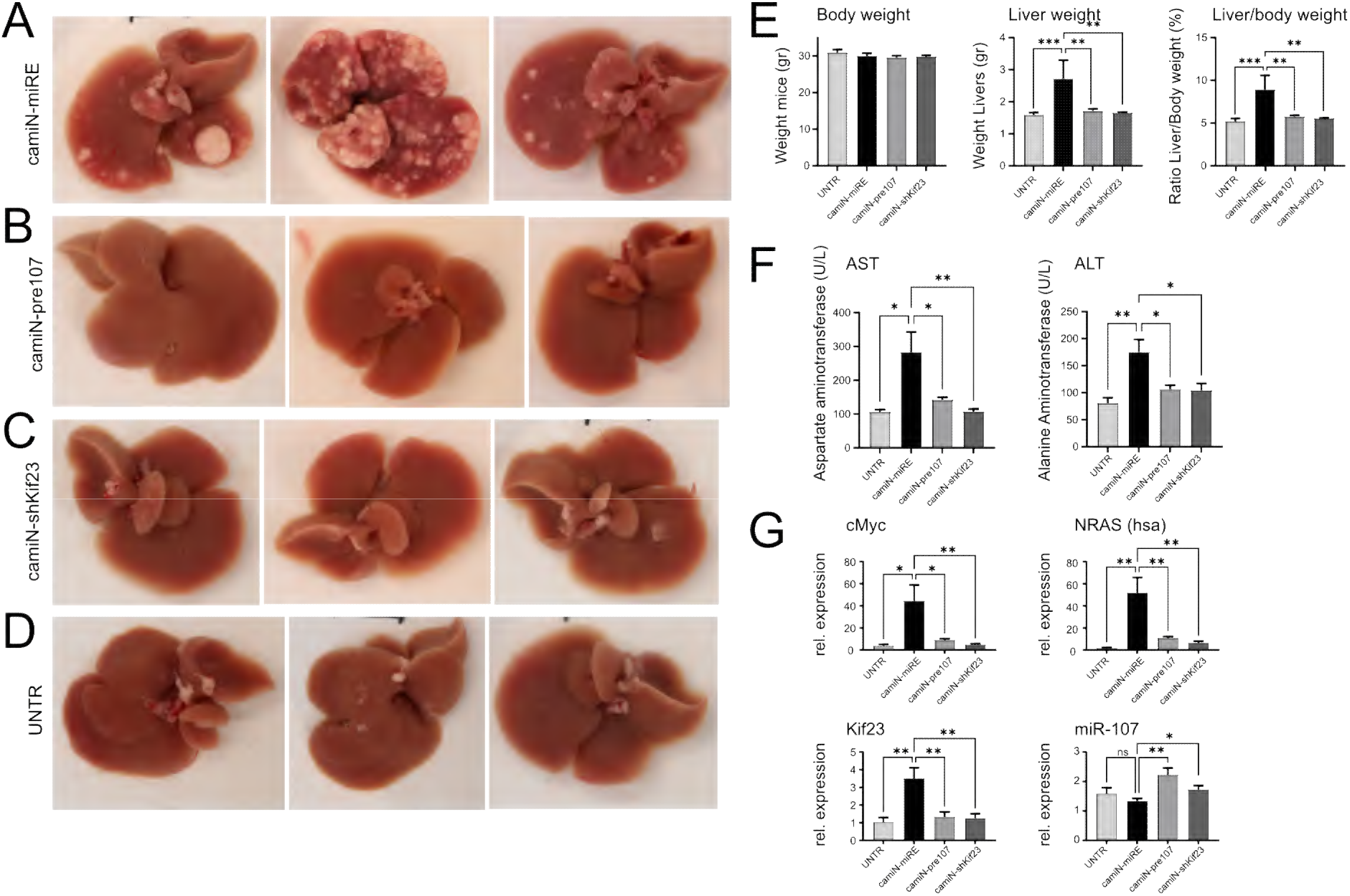
Targeting the miR-107/Kif23 axis prevents the formation of oncogene- induced liver cancer. Representative images of tumor burden six weeks after HDTVi- mediated administration of transposonal vectors carrying the cMyc/NRas oncogenes in combination with either (**A**) the control miRNA (camiN-miRE), (**B**) the pre-miR-107 (camiN-pre-107), (**C**) the anti-Kif23 siRNA (camiN-shKif23), (**D**) or left untreated (UNTR). The effect of oncogene administration in combination with miR-107 overexpression and Kif23 inhibition on liver cancer development was evaluated at physiological level by measuring (**E**) body and liver weight parameters and (**F**) liver enzymes as shown for AST and ALT and at molecular level by RT-qPCR quantification of the oncogenes (**G**) cMyc and NRAS (hsa) as well as for the endogenous Nras (mmu) and by the quantification of Kif23 and miR-107 expression. UNTR (n = 4) or HDTVi- injected (n = 7 per treatment group) mice. Results are represented as mean ± SD, significant differences were evaluated by using 1-way ANOVA with Newman-Keuls post-hoc test. (ns = Not significant, *, p ≤ 0.05; **, p ≤ 0.01; ***, p ≤ 0.001).

### Kif23 inhibition, but not miR-107 overexpression, completely prevented liver tumor formation

To determine the extent to which miR-107 overexpression and Kif23 silencing could protect the liver from oncogene-induced malignant transformation, we conducted a histological analysis of the livers of these animals to exclude or identify the presence of abnormal cellular features and to investigate the possible development of malignancy at the microscopic level. Consistent with the macroscopic analysis of the livers, serum markers, and molecular markers, the histopathological analysis of hematoxylin and eosin (H&E)-stained formalin-fixed, paraffin-embedded (FFPE) liver slices from shKif23-treated animals revealed no abnormalities associated with cellular dysplasia, nodules, or oncogenic transformation (**Figure 7, Supplementary Table 7**). In contrast to the macroscopic evaluation, molecular marker analysis, and serum marker analysis, histopathological analysis of the livers of animals injected with vectors encoding pre-miR-107 and the cMyc/NRas oncogenes revealed diffuse hepatocellular carcinoma characterized by small, well-defined nodules (**Figure 7C**).

**Figure 7.**
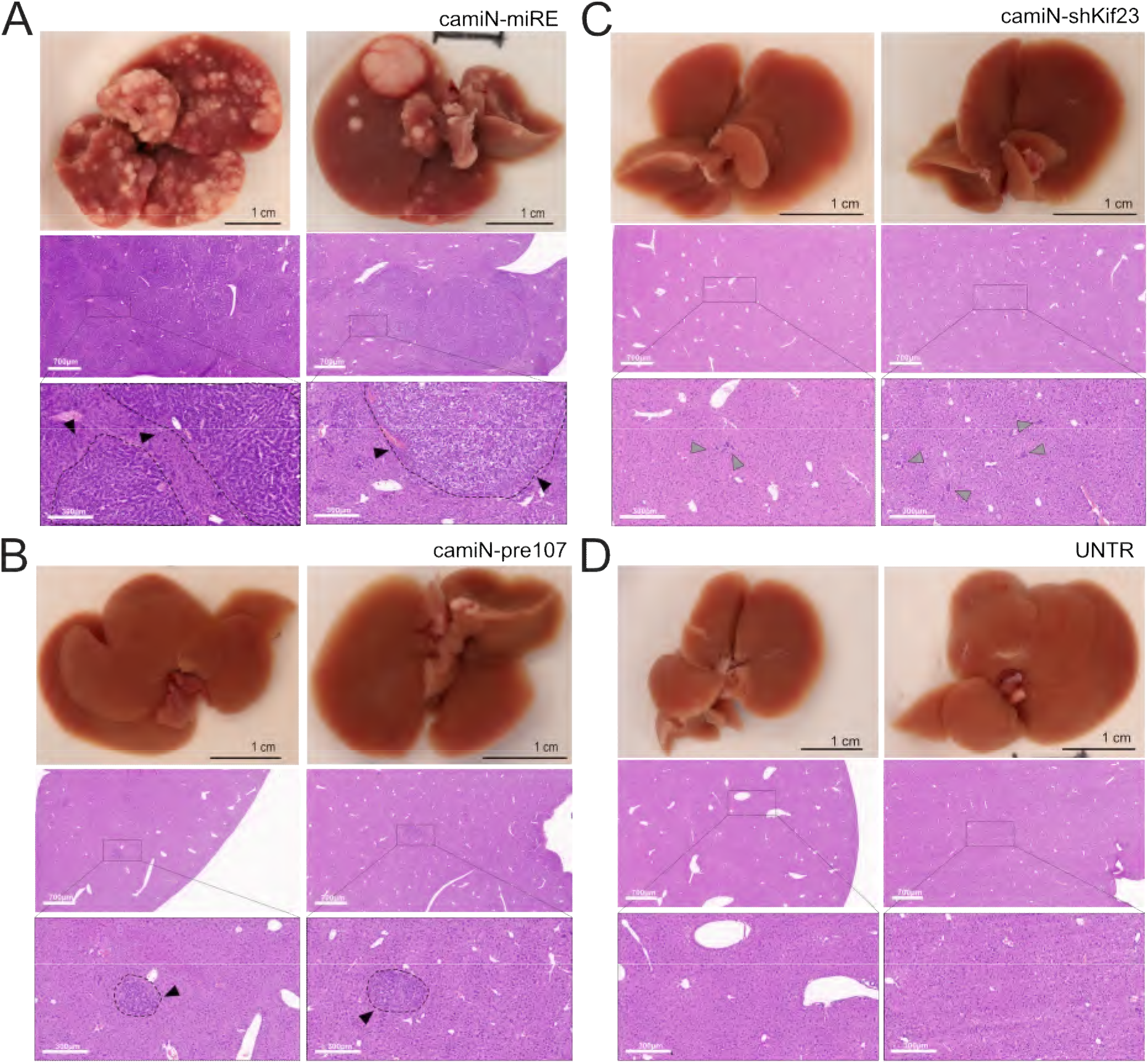
shRNA-mediated inhibition of Kif23 prevents the formation of liver cancer, while miR-107 overexpression delays cancer development but does not prevent it entirely. Representative images showing hematoxylin and eosin (HE)- stained slices from FFPE blocks prepared from the livers of wild-type mice six weeks after the HDTVi-mediated administration of transposonal vectors carrying the cMyc/NRas oncogenes administered in combination with either the (**A**) control miRNA (camiN-miRE), (**B**) the pre-miR-107 (camiN-pre-107), (**C**) the anti-Kif23 siRNA (camiN- shKif23), or (**D**) they were left untreated (UNTR). Tumors can be observed in the livers of animals injected with camiN-miRE and in the camiN-pre107 vectors (black arrow), whereas the livers of camiN-shKif23 and UNTR animals remained free from liver cancer. In the camiN-shKif23 injected animals small number of atypical cells were detected (gray arrow).

These findings were further corroborated by the analysis of the immune cell landscape in the livers of these animals. To assess the distribution and quantity of immune cells, liver sections were stained with F4/80 and CD3 to visualize macrophages and T cells, respectively. Notably, the spatial distribution and number of T cells remained essentially unchanged (**Supplementary** Figure 6), while the macrophage population was significantly reduced in the livers of both groups of animals injected with miRE and pre-miR-107, compared to the macrophages in the livers of untreated animals. Importantly, no significant differences in the degree of macrophage infiltration were observed between the livers of untreated control animals and shKif23-injected animals (**Supplementary** Figure 7).

Overall, these findings support the conclusion that shRNA-mediated silencing of Kif23 protects the liver from highly aggressive oncogene-induced HCC, confirming our initial hypothesis that Kif23 is required for supporting the fitness of proliferating cancer cells. Thus, Kif23 represents a promising therapeutic target for hepatocellular carcinoma.

## Conclusion

Hepatocellular carcinoma, the most common form of liver cancer, is characterized by a high incidence, limited treatment options, and a high mortality rate. The global incidence of HCC is anticipated to rise in the coming years, primarily due to the ongoing epidemic of obesity, which is a leading risk factor for HCC. The lethality of HCC and the limited therapeutic options available highlight the urgent need to identify new targets for treatment. Despite significant advances in cancer research, the translation of preclinical research from animal studies to clinical trials remains a challenge, particularly in the context of liver cancer. Animal models commonly used to study liver cancer do not fully replicate the heterogeneity observed in humans [36], thus hindering the successful translation of preclinical studies into fully fledged clinical trials [37].

Over the past decade, miRNAs have emerged as key regulators of gene expression in cancer, including liver cancer. Notably, the dysregulation of miRNAs expression, along with their associated target genes and regulatory signaling pathways, has been shown to be conserved across species, as evidenced by a growing body of literature [38–40]. To overcome these challenges, we developed an innovative strategy to globally identify miRNAs and their target genes that undergo significant alterations in liver cancer [6]. This approach successfully revealed miRNA-driven transcriptional networks associated with liver cancer in mice, which were found to be similarly dysregulated in the livers of HCC patients. In the present study, the role of miR-107 in hepatocarcinogenesis was explored. The deregulation of miR-107 expression has been implicated in various human cancers, with its role as either a tumor suppressor or an oncogene being debated [41–43]. This debate also extends to liver cancer, where studies have reported both carcinogenic [44] and tumor-suppressive effects of miR- 107 [45, 46].

Through extensive analysis, we demonstrated that downregulation of miR-107 is a hallmark of liver cancer in mice. We also found that lower miR-107 expression correlated with enhanced fitness of human and mouse hepatoma cells and significantly lower survival in LIHC patients. These findings support the conclusion that miR-107 is a tumor suppressor miRNA and that its downregulation promotes liver cancer development. Based on our data and the current literature, it can be proposed that miR-107 might act as a double-edged sword displaying both anti- and pro-tumorigenic activity, that might be regulated by distinct oncogenic and tumor suppressor circuits. Based on this hypothesis, identical changes in miR-107 expression may lead to different outcomes depending on the upstream regulator or cancer type, in a tissue- and cell-specific manner. miR-107 is a member of the miRtron family, as the sequence encoding for miR-107 primary transcript is located in an intron of the Pantothenate kinase 1 (Pank1) gene [41]. Interestingly, miR-107 expression does not directly correlate with Pank1 transcription [47], suggesting that miR-107 biogenesis and processing are regulated by different signaling pathways than Pank1. Elucidating the mechanisms underlying miR-107 regulation could have significant implications for understanding the regulatory networks and role of miR-107 in a specific cancer. Notably, miR-107 activity was shown to play a role in the regulation of a number of signaling pathways, including PTEN/AKT [48], PI3K/Akt [49], STK33/ERK [50], and TGF-β signaling [51], suggesting that miR-107 may act as a pleiotropic regulator in liver cancer development.

To better understand the role of miR-107 as a tumor suppressor gene in liver cancer, we sought to identify miR-107-regulated genes that act as functional mediators of its activity and supporting the fitness of cancer cells. Through comprehensive approaches combining system biology and experimental validation, we identified KIF23, a spindle-associated microtubule motor protein, as a novel direct target of miR-107. KIF23 is a key component of the centralspindlin complex, which is required for central spindle assembly and cytokinesis [52]. KIF23 plays a crucial role in cell division, particularly during mitosis, where it is required for the formation of the contractile ring and the subsequent cleavage of the cell membrane of the two daughter cells. Regarding its potential role in cancer, elevated KIF23 levels have been observed in a number of human cancers, including hepatoma [26], colorectal cancer [23], pancreatic ductal adenocarcinoma [24], and ovarian cancers [25], where high levels of KIF23 have been associated with poor prognosis. Although the precise mechanisms through which KIF23 influences cancer development and progression are not fully understood, it is known that KIF23 per se doesn’t promote tumorigenesis or oncogenic transformation. Therefore, we propose that KIF23 enhances tumor cell fitness by supporting cellular division, which indirectly promotes tumor cell proliferation. Through our work, we showed that high expression levels of KIF23 were associated with the malignant behavior of human hepatoma cells and correlated with unfavorable clinical outcomes in cohorts of LIHC patients. Notably, through the analysis of TMA prepared from the liver cores of HCC patients, we could show that KIF23 protein accumulates in the nuclei of tumor cells compared to the nuclei of cells localized in the surrounding, non-tumorigenic tissue.

In recent years, methods to inactivate spindle-associated kinesins have been extensively investigated because silencing specific members of this class of motor proteins would provide an effective therapy against cancer. However, despite promising *in vitro* data, the introduction in clinical trials of inhibitors targeting this category of proteins, including “ispinesib” targeting the Kinesin spindle protein (KSP/Eg5) and “GSK923295” targeting the Centromere protein E (CENP-E), failed to meet their initial promise [53–55]. These setbacks highlight the need to explore alternative approaches and methods for targeting this category of proteins. In this respect, our discovery that siRNA-mediated silencing of Kif23 prevents cMyc/NRas- induced liver cancer formation *in vivo* qualifies Kif23 as a novel promising target. It is interesting to highlight that despite the complete absence of macroscopic and microscopic tumors, some small abnormalities were detected following the histological analysis of the H&E staining of livers from the mice injected with the vector encoding for the oncogenes and anti-Kif23 shRNAs. We propose that these abnormalities are dysplastic foci containing a small number of atypical cells, indicating that the oncogene-induced neoplastic process in the cells receiving the transposon vectors was halted early, hence preventing the formation of full-fledged lesions. We speculate that these dysplastic foci might be linked to the immune-mediated clearance of pre- malignant senescent parenchyma cells. While the silencing of KIF23 resulted in a black-and-white phenotype, the same cannot be said about the overexpression of its regulatory gene, miR-107. Overexpression of miR-107 did not lead to a tumor-free phenotype, which raises intriguing questions. To answer these questions, we propose two hypotheses: (i) miR-107 is a tumor suppressor, but its overexpression is not sufficient to fully suppress tumor growth, and (ii) miR-107 is a double-edged sword, with both tumor-suppressive and tumor-promoting functions. In support of the first hypothesis, miRNAs are known to modulate, not inhibit, their target genes, hence it is possible the overexpression of miR-107 is not sufficient to fully suppress Kif23 and to block tumor growth, hence resulting in the observed delay in tumor formation. In support of the second hypothesis, miR-107 targets several other genes besides KIF23. Therefore, while its upregulation may initially protect against cancer development, over time, the synergistic effects of miR-107 suppression of other targets/signaling pathways could lead to malignancy. To this end, the observation that human NRas, one of the oncogenes introduced with HDTVi, was significantly increased in the livers of the miRE but not in the livers of shKif23 and pre-miR-107 animals would support this conclusion.

While this finding does not “disqualify” miR-107 from being considered a therapeutic target, previous studies have suggested that interfering with miR-107 *in vivo* may carry risks. Based on the discovery that miR-107 regulates insulin sensitivity [56], AstraZeneca/Regulus developed RG-125 (AZD4076), an RNA therapeutic targeting miR-107 *in vivo*. The efficacy of this therapeutic was assessed in clinical trials involving patients with type 2 diabetes and Metabolic dysfunction-associated steatohepatitis (MASH; the replacement term for NASH). However, the study was terminated in 2017. Interestingly, an independent study demonstrated that targeting miR-103/-107 had adverse effects on cardiac function and metabolism [57]. Although we cannot definitively establish a causal relationship between these two events, the observation suggests that systemic targeting of miR-107 could potentially have detrimental effects on human health. This, coupled with the demonstrated role of miR-107 downregulation in liver cancer development, necessitates a cautious re-evaluation of systemic miR-107 targeting specifically and miRNA targeting in general.

While our study has identified Kif23 as a promising new therapeutic target in HCC, its limitations must be acknowledged. First, it is not clear whether targeting Kif23 can induce regression of liver cancer once it has become established. Second, the HDTVi method used in this study is not suitable for delivering RNA therapeutics to humans. Therefore, alternative delivery strategies, such as lipid nanoparticles, engineered extracellular vesicles, or cholesterol-conjugated siRNA, should be investigated for their efficacy in delivering RNA therapeutic targeting Kif23 *in vivo*.

In summary, our work contributes to our understanding of the molecular mechanisms involved in liver cancer development, as summarized in the model outlined in **Figure 8**. By discovering and exploring the role of the miR-107/KIF23 axis in mouse liver cancer, we shed light on a novel potential target for liver cancer in humans. This may pave the way for the development of novel, more effective therapeutic strategies in our fight against this deadly disease.

**Figure 8.**
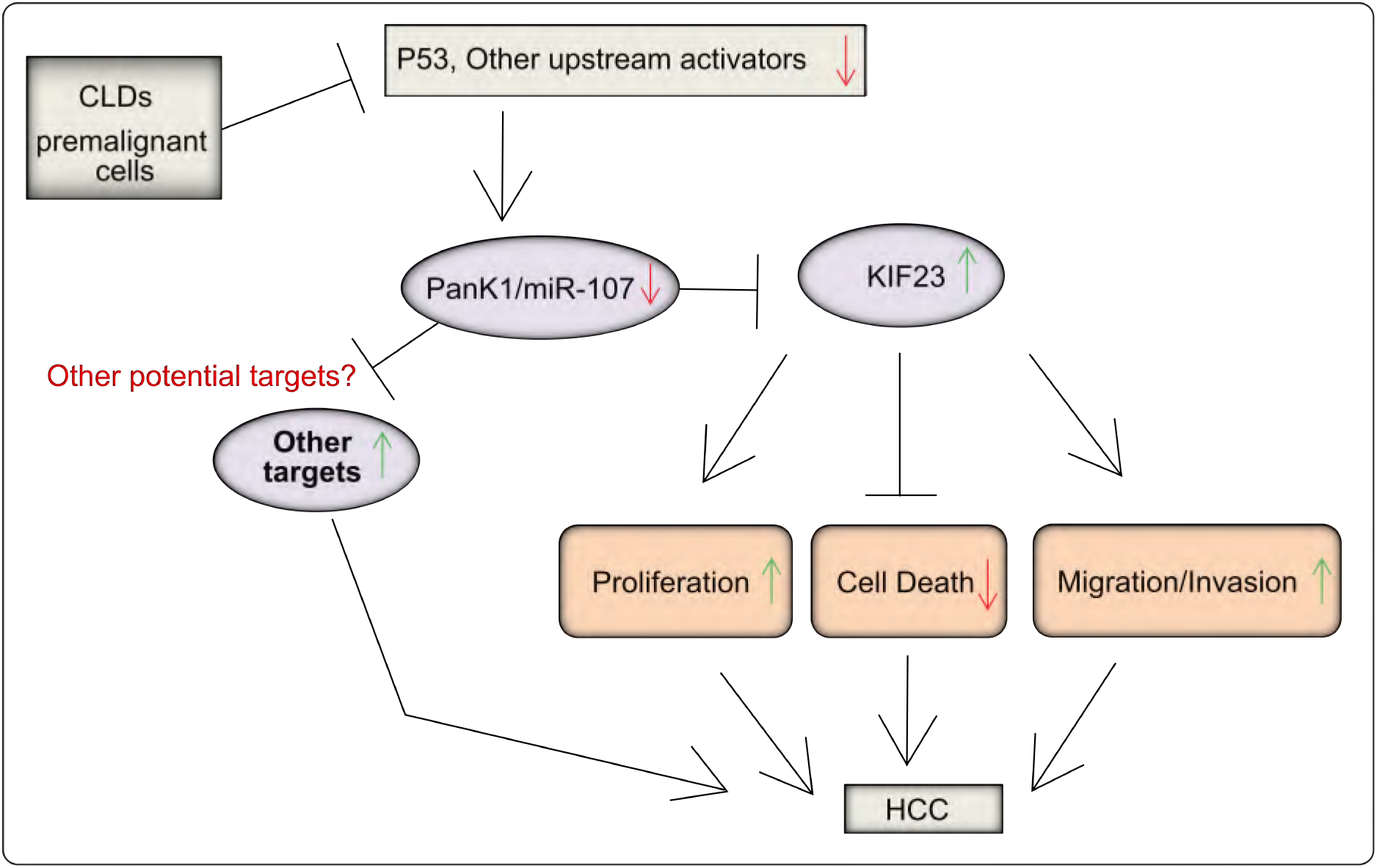
Proposed model for the regulation of the miR-107/Kif23 axis in liver cancer. We propose that miR-107 levels are downregulated in chronic liver diseases (CLDs) and in pre- and malignant cells due to the deregulation or activation of yet-to-be-identified signaling pathways that modulate miR-107 biogenesis. This downregulation leads to an upregulation of miR-107 targets, including KIF23. While KIF23 is not classified as an oncogene, it supports cell division by driving cytokinesis. This enhances the fitness of cancer cells and results in a significant increase in proliferation and migration, indirectly reducing cell death.

## Abbreviations

miRNA, microRNA; ncRNAs, noncoding RNAs; shRNA, short hairpin RNAs; siRNA, short interfering RNA; 3’ UTRs, 3’ untranslated regions; MRE, miRNA responsive elements; GSEA, Gene set enrichment analysis; HCC, hepatocellular carcinoma; NTL, near-tumor liver; HDTVi, Hydrodynamic tail vein injection; TMAs, Tissue Microarrays; MASH/(NASH), Metabolic dysfunction-associated steatohepatitis; CLDs, Chronic liver diseases; TCGA, The Cancer Genome Atlas; GEO, Genome Omnibus; Kif23, Kinesin family member 23; Aurkb, Aurora Kinase B; Prc1, Polycomb repressive complex 1; Ttk, Monopolar spindle 1; Anln, Anillin; Nuf2, Nuclear Filament-containing protein 2; Pank1, of Pantothenate kinase 1; KSP/Eg5, Kinesin spindle protein; CENP-E, Centromere protein E; DEN, Diethylnitrosamine; LIHC, liver hepatocellular carcinoma cohorts; TUNEL, Terminal deoxynucleotidyl transferase dUTP nick end labeling; ALT, Alanine transaminase; AST, Aspartate Transferase; GLDH, Glutamatdehydrogenase; HE, hematoxylin and eosin; IHC, Immunohistochemistry; IF, Immunofluorescence.

## Acknowledgements

The authors are grateful to Claudia Rupprecht, Nicole Eichhorst, Stefanie Lindner, Michaela Fastrich, Torsten Janssen and Nathalie Walter for the technical assistance. The authors thank the Center of Model System and Comparative Pathology (CMCP) at the Institute of Pathology in Heidelberg for their technical assistance in human tissue staining and the Tissue Bank of the National Center of Tumor Diseases (NCT, Heidelberg) for the tissue slide scanning.

## Author Contributions

M.C. and T.Lu., designed and guided the research. M.C. and T.Lu., wrote the manuscript with help from other authors. M.C., performed and analyzed most of the experiments. S.R., C.L., R.P., M.V., M.t.S., V.D., M.a.D, and C.R., contributed to research design and/or conducted experiments. L.Z., provided important technical support. R.P., L.r.H, U.p.N., and T.Lo., generated and analysed the Tissue microarrays. T.Lo., conducted histo-pathological analyses.

All the authors read and approved the manuscript.

