## Supplementary Figures for "KIF23 regulation by miR-107 controls replicative tumor cell fitness in mouse and human hepatocellular carcinoma"

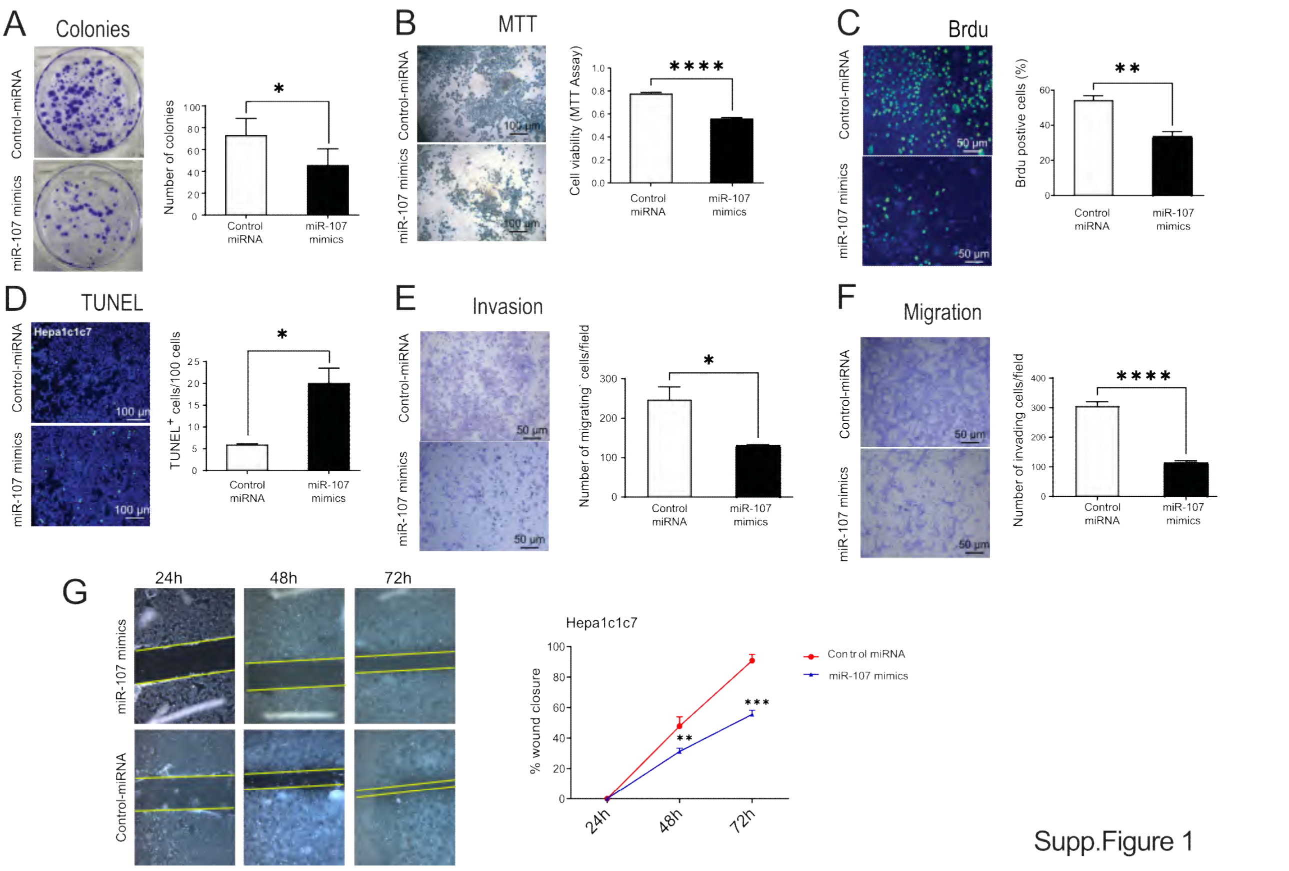


**Supplementary Figure 1. miR-107 upregulation, negatively impact the fitness of mouse hepatoma cells.** To assess the effect of miR-107 on the hallmark of cancer mouse-derived hepatoma Hepa1c1c7 were transfected with either miR-107 mimic or control miRNA, and the effect on the fitness of cancer cells was measured by using: (**A**) colony formation assay. (**B**) MTT proliferation assay (n = 5), (**C**) BrdU proliferation assay (n = 5), (**D**) cell viability was measured by TUNEL assay (n = 5). Transwell migration assay (**E**) or Transwell coated with Matrigel invasion assay (**F**) for Hepa1c1c7 cells was determined after transfection with miR-107 mimic or control miRNAs for 72 h (n = 3). (**G**) Wound-healing assay was performed on Hepa1c1c7 cells transfected with miR-107 mimic and wound closure was monitored over the indicated time post-transfection after treatment with Mitomycin C for 2 h (n = 3). Representative images are shown for the different experiments. Results are represented as mean ± SD, significant differences were evaluated by 2- tailed, unpaired t test t-test as appropriate. (*, $p \leq 0.05$; **, $p \leq 0.01$; ***, $p \leq 0.001$; ****, $p \leq 0.0001$).

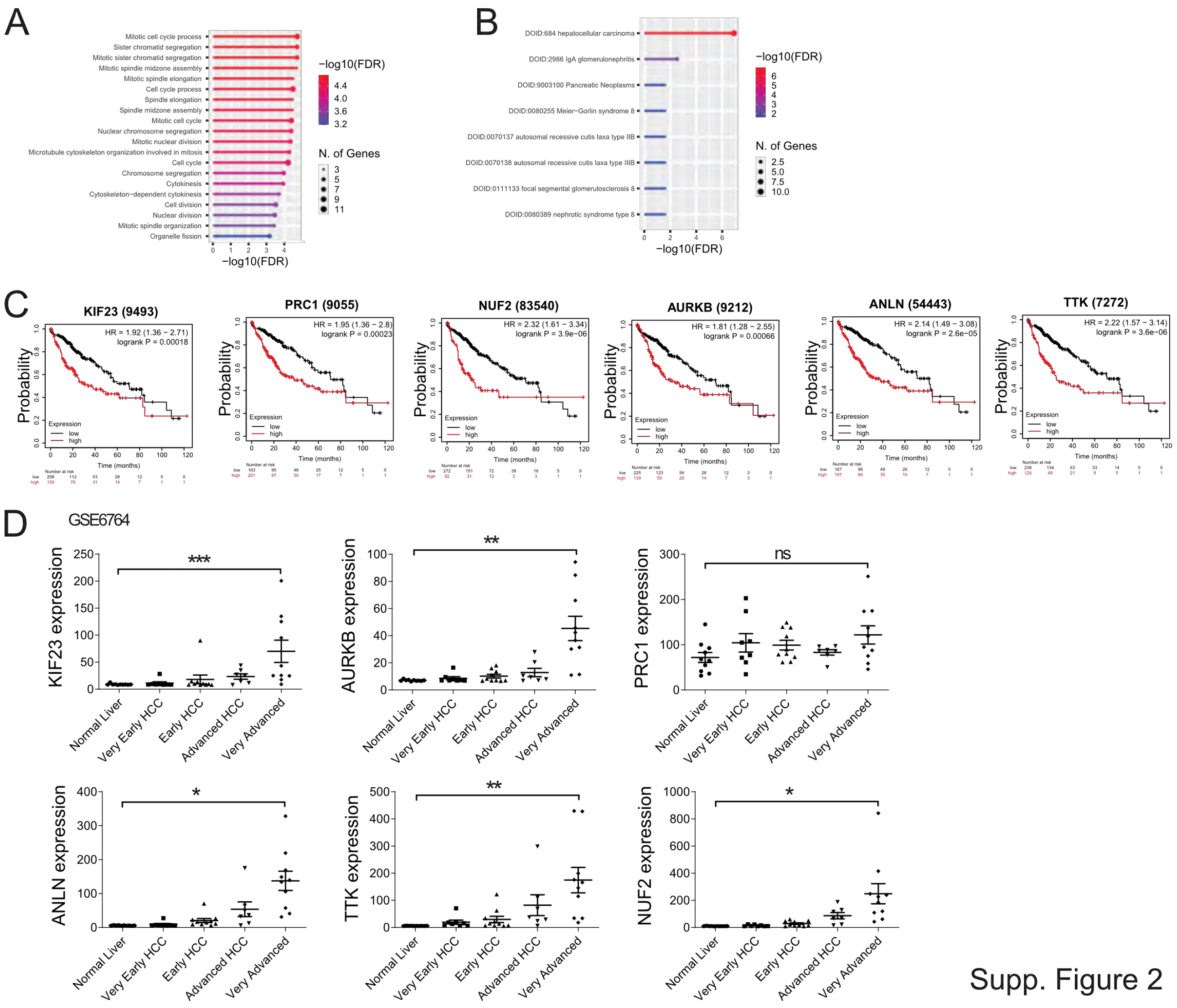


**Supplementary Figure 2. miR-107 predicted targets in mouse; a potential role in human HCC.** (**A**) ShinyGO enrichment analysis of the 23 potential miR-107 targets and (**B**) and ShinyGO ebrichment analysis for diseases using the “Disease:Jensen:Diseases” database. (**C**) Kaplan-Meier survival curves for KIF23, PRC1, NUF2, AURKB, ANLN and YYK, in TCGA LIHC cohorts. (**D**) The TCGA data were independently validated via the analysis of the GSE6764 cohort that contains the comparison between tumor tissues at different stage.

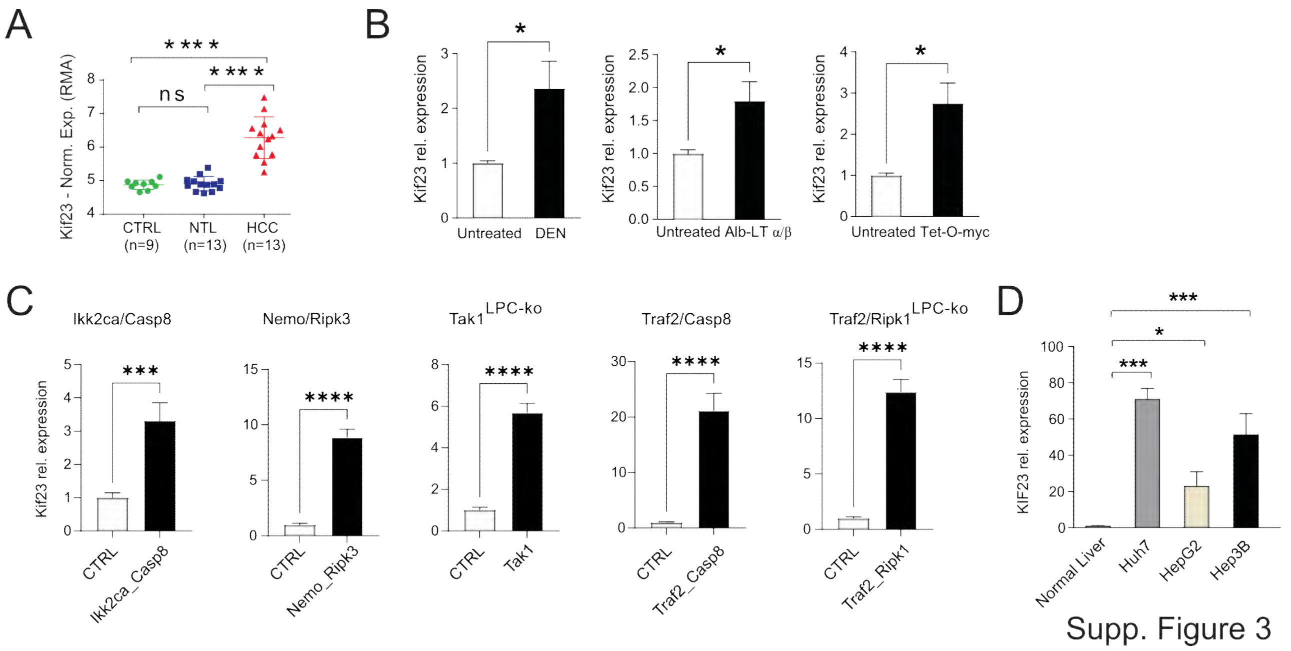


**Supplementary Figure 3. Kif23 is significantly down-regulated in mouse models of different etiology and in human hepatoma cell lines.** (**A**) RMA-normalized microarray data for the expression of Kif23 in the healthy (n = 9), nontumoral liver (NTL: n = 13) and tumor (n = 13) liver tissue from the mice included in this study. (**B**) RT-qPCR validation of Kif23 expression in the individual DEN (Untreated, n = 6; Tumor, n = 5), AlbLTα/β (Untreated, n = 4; Tumor, n = 4), and Tet-O-Myc (Untreated, n = 4; Tumor, n = 4) models of liver cancer. (**C**) RT-qPCR analysis of Kif23 expression in five different genetic HCC mouse tumor models: Ikk2/Casp8 (n = 5), Nemo/Ripk3 (n = 5), Tak1$^{LPC\text{-}ko}$ (n = 5), Traf2/Casp8$^{LPC\text{-}ko}$ (n = 5), and Traf2/Ripk1 $^{LPC\text{-}ko}$ (n = 5) compared to control (CTRL; n = 5) mice. (**D**) RT-qPCR of KIF2 3 expression in normal human (n = 3) livers and in and human hepatoma cells (Huh-76; n = 5, HepG2; n = 5, Hep3B; n = 5). Results are represented as mean ± SD, significant differences were evaluated by using 1-way ANOVA with Newman-Keuls post-hoc test or 2- tailed, unpaired t test t-test as appropriate. (ns = Not significant, *, $p \leq 0.05$; **, $p \leq 0.01$; ****, $p \leq 0.0001$).

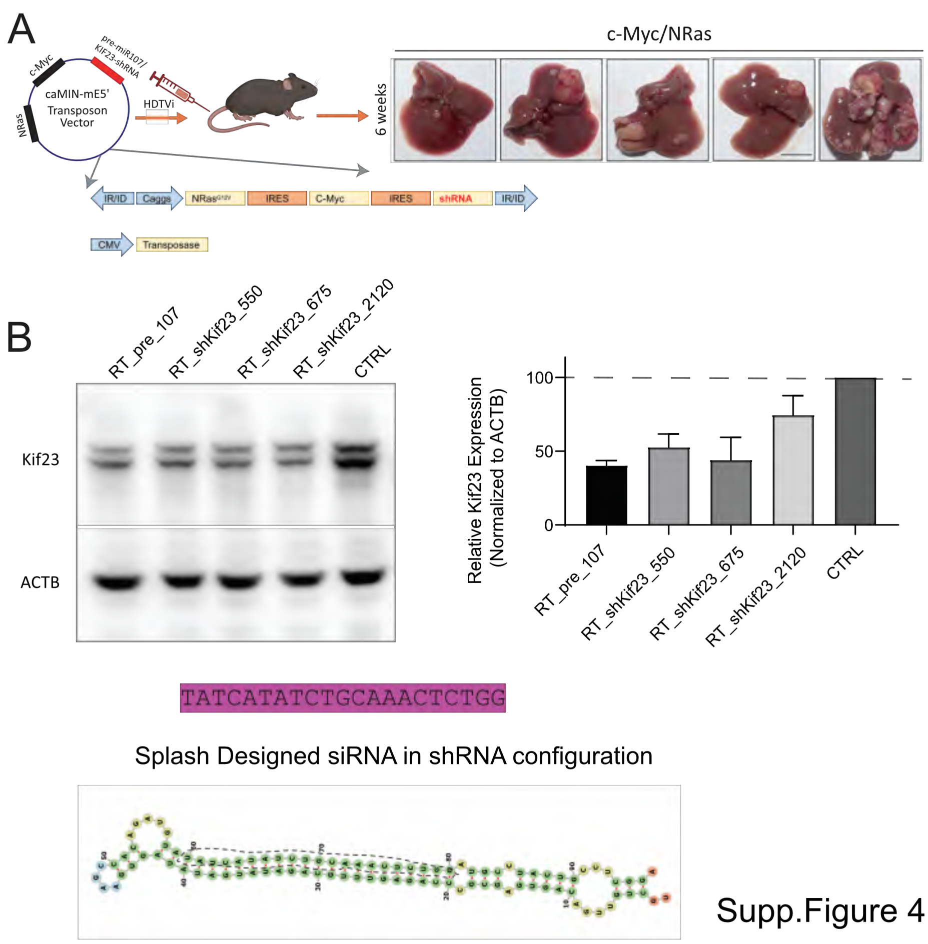


Supp.Figure 4

**Supplementary Figure 4. Design, strategy and validation of shRNA targeting Kif23 for HDTVi delivery.** (**A**) The design of siRNA, cloning and subcloning in the transposon vector encoding for cMyc/NRas oncogenes has been previously described (REF). The only difference between is that the published method used a three vectors system for delivering, oncogene, shRNA and transposase. In our study we used a simplified two vectors system (**B**) Three different siRNA where designed and cloned in the entry vector 3RT-GEPIR. These vectors, along with a vector carrying the sequence 1 encoding for pre-miR-107 where transiently transfected in mouse Hepa1-6 cells. Three days after transfections cells were lysed, and Kif23 expression was evaluated by WB. The most efficient siRNA (RT_shKif23_675), along with the RT_pre_107 were subcloned in the camiNm5E vector and used for the HDTVi. The secondary structure of the shKif23_675 is shown in the lower part of the figure. Results are represented as mean ± SD

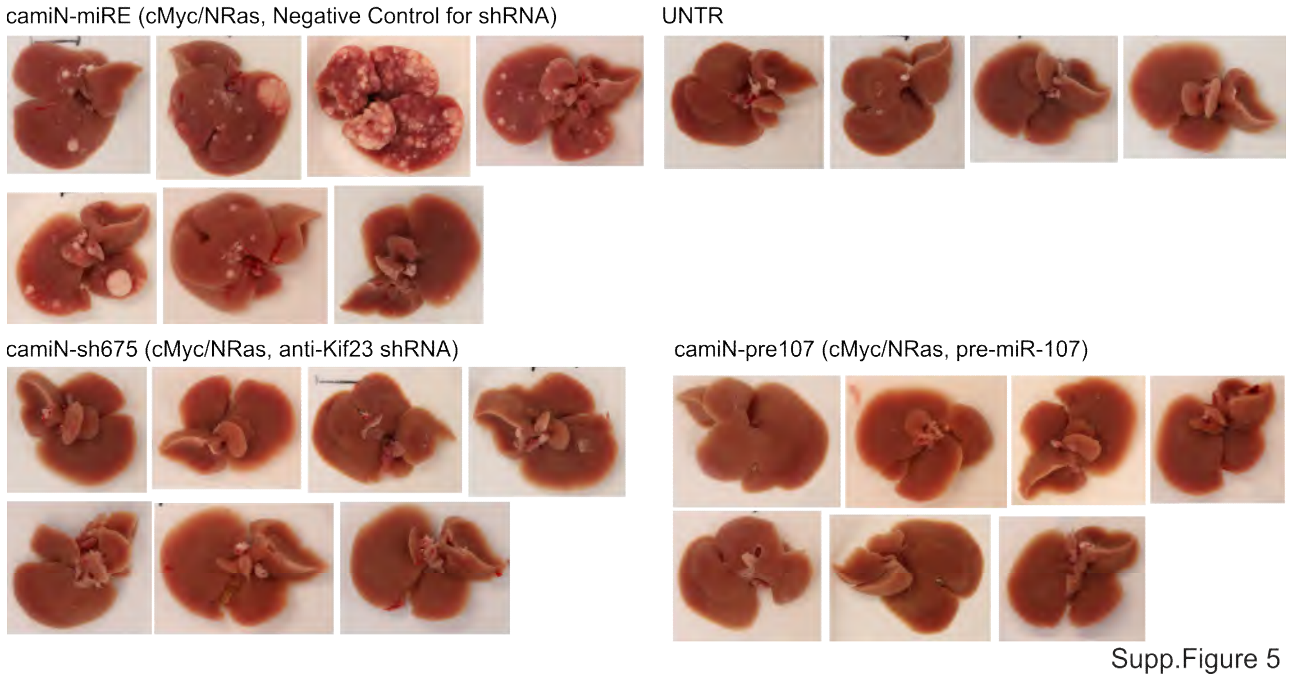


**Supplementary Figure 5. Complete overview of the liver from the HDTV injected mice.** Tumor burden six weeks after HDTVi-mediated administration of transposonal vectors carrying the cMyc/NRas oncogenes in combination with either the control miRNA (camiN-miRE), the pre-miR-107 (camiN-pre-107), the anti-Kif23 siRNA (camiN-shKif23), or left untreated (UNTR).

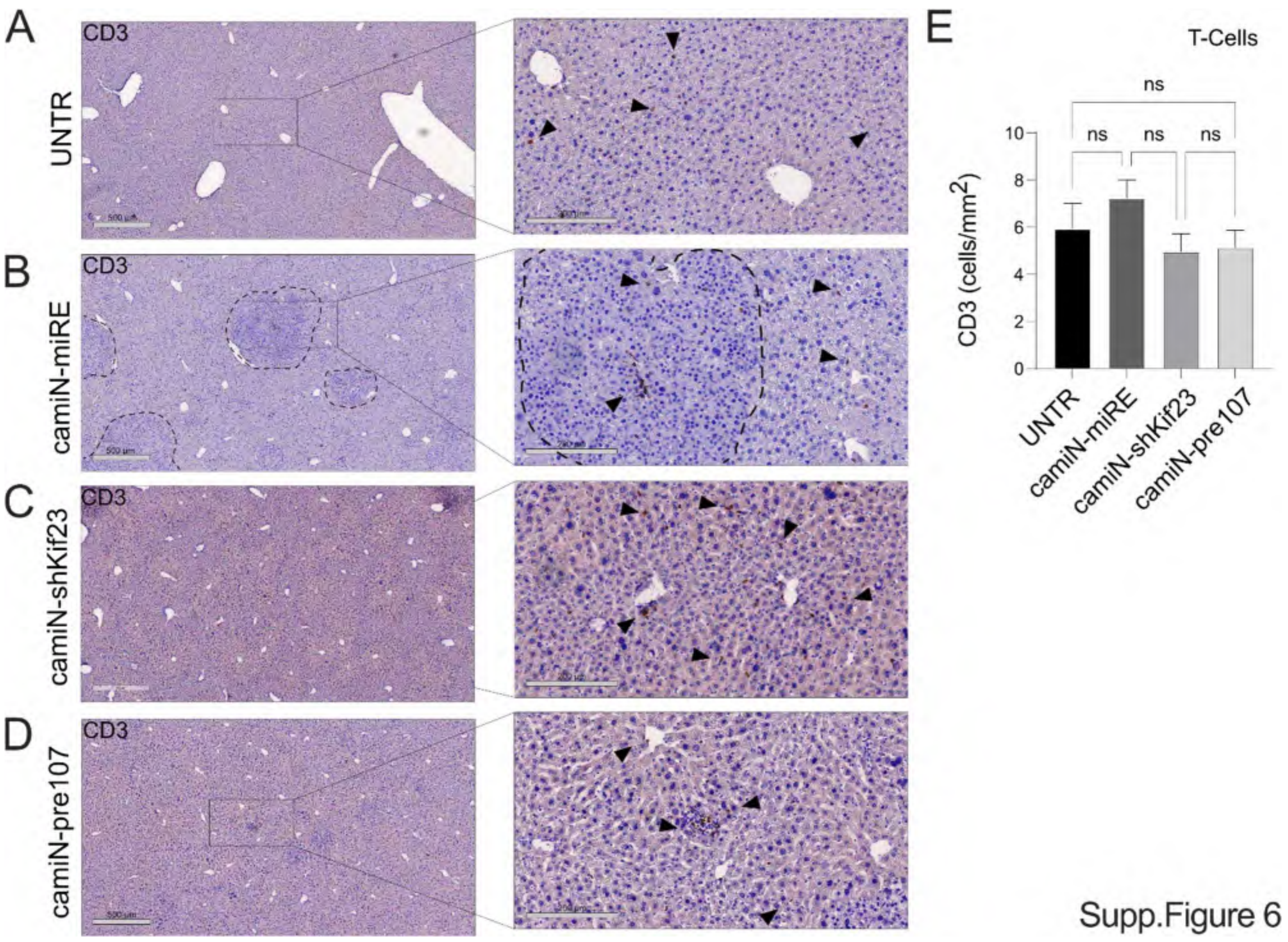


**Supplementary Figure 6. CD3 staining of liver tissue from HDTVi injected and control mice.** CD3 staining did not show altered inflammation/distribution pattern in the livers of any of the animals included in the study. (**A**) Untreated (UNTR) animals, (**B**) control miRNA (camiN-miRE), (**C**) the anti-Kif23 siRNA (camiN-shKif23) and (**D**) the pre-miR-107 (camiN-pre-107) injected animals. (**E**) Quantification of CD3 positive cells. Results are represented as mean ± SD, significant differences were evaluated by using 2- tailed, unpaired t test (ns; Not Significant).

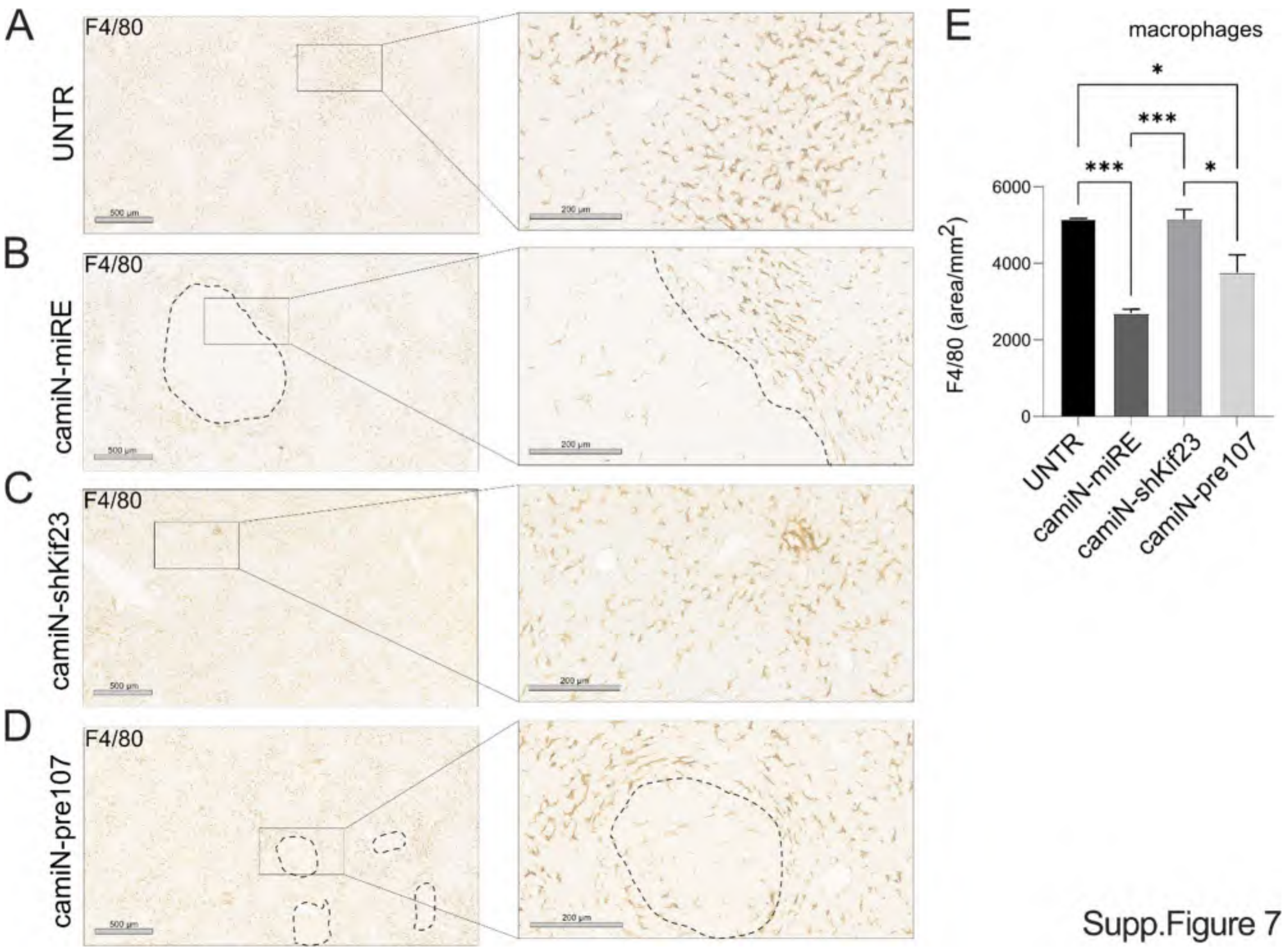


**Supplementary Figure 7. F4/80 staining of liver tissue from HDTVi injected and control mice.** F4/80 staining did not show altered inflammation the liver of (**A**) UNTR and (**C**) mice injected with the transposonal vectors carrying the cMyc/NRas oncogenes anti-Kif23 siRNA (camiN-shKif23). A significant reduction on the number of macrophages was detected in the livers of mice six weeks after the HDTVi-mediated administration of transposonal vectors carrying the (**B**) cMyc/NRas oncogenes in combination with either the control miRNA (camiN-miRE), (**D**) the pre-miR-107 (camiN-pre-107). The left panel show representative images at different magnification. (**E**) Quantification of F4/80 signals. Results are represented as mean ± SD, significant differences were evaluated by using 2- tailed, unpaired t test (*, $p \leq 0.05$; ***, $p \leq 0.001$).

### Legends for the Supplementary Tables

**Supplementary Table 1**

Clinical characteristics of patients who donated their liver tissues for the preparation of the tissue microarrays (TMAs).

**Supplementary Table 2**

Differentially expressed genes in DEN, AlbLTα/β, and Tet-O-Myc livers compared to controls. Data from the GSE102417 dataset were downloaded from the Gene Expression Omnibus (GEO) repository. Affymetrix CEL files were analyzed with AltAnalyze (version 2.1.4.3). Genes were considered differentially expressed if they met both of the following criteria: Fold change (FC) ≥ 2.0 relative to the control group. Statistical significance at $p \leq 0.05$. For the purpose of increased specificity, the CEL files were divided into three groups: Control animals (CTRL), Non-tumor liver (NTL): DEN, AlbLTα/β, and Tet-O-Myc livers with minimal to no tumor burden, Tumor liver: DEN, AlbLTα/β, and Tet-O-Myc livers with a high tumor burden

**Supplementary Table 3**

List of predicted miR-107 responsive genes, obtained by overlapping the gene list in Supplementary Table 2 with the complete list of predicted miR-107 targets in the miRWalk target prediction database.

**Supplementary Table 4**

Kif23 immunohistochemistry (IHC) scores for tumor (T) and matching non-neoplastic (N) liver tissue in Tissue Microarray (TMA) prepared from human livers (n = 62).

**Supplementary Table 5**

Measurement of body weight, liver weight, and the liver weight-to-body weight ratio in 13-week-old C57BL/6J male mice subjected to different experimental conditions 6 weeks after HDTVi. Data are expressed as means ± SD.

**Supplementary Table 6**

Levels of serum markers (AST, ALT, ALP and GLDH) measure in the blood of the mice included in this study 6 weeks after HDTVi. HDTVi were performed on 7 weeks old C57BL/6J male mice. Animals were euthanized 6 weeks after HDTVi. Data are expressed as means ± SD

**Supplementary Table 7**

Pathological analysis of the H&E stained slides of the livers of the mice included in this study. morphological hepatocarcinogenesis start from dysplastic foci, which than develop to dysplastic nodule (>1 mm diameter). HCC develops from the latter lesion (and sometimes borderline cases between DN and HCC can be observed).

**Supplementary Table 8**

Sequences of oligonucleotides used for miRNA and mRNA expression profiling by RT-qPCR, as well as oligonucleotide sequences for cloning of human and mouse KIF23 3' UTRs in the pMIR luciferase vector and sequence of the oligonucleotides used to lone the KIF23-shRNA and pre-miR-107 constructs in the RT3GEPIR and camiNmA vectors. Sequences are reported in the 5'-to-3' orientation.

**Supplementary Table 9**

List of miRNA mimics/inhibitors and siRNAs used in this study
