## Supplementary Material and Methods for "KIF23 regulation by miR-107 controls replicative tumor cell fitness in mouse and human hepatocellular carcinoma"

### Supplementary Materials and Methods

#### RNA Isolation, Quantification, cDNA Synthesis, and qPCR Analysis

RNA isolation, quantification, and reverse transcription were performed as previously described [1]. miRNA profiling was performed by using the miQPCR method [2]. miQPCR performs the universal reverse transcription of all the miRNAs contained in the RNA sample enabling the analysis of up to 100 individual miRNAs from every synthesized cDNA. Amplicon amplifications were carried out on a ViiA7 cycler (Thermo Fisher Scientific, Meerbusch, Germany) and detected using SYBR Green I (GO-Taq PCR Master mix, Cat: A6002; Promega, Walldorf, Germany). Sequences of the primers used in this study are listed in **Supplementary Table 8**. Data analysis was carried out using qBase [3]. Suitable reference genes were identified using geNorm [4], and samples were normalized by using ΔΔCt [5]. GraphPad Prism (GraphPad Software, Inc., La Jolla, CA, Version 9.4.1) was used to perform statistical analyses and for data visualization.

#### MTT, BrdU, TUNEL and colony formation assays.

To measure viability, cells were seeded in 24-well plates and transfected with miR-107 mimics or Kif23-siRNAs or scrambled oligos and let recover for 72 h. Following cells were either incubated with MTT solution (Vybrant MTT cell proliferation assay, Molecular Probes, Eugene, OR, USA, V-13154) or for TUNEL assay (Promega, Walldorf, Germany, G3250), and processed according to the provided protocol. Absorbance was measured in 540 nm by Biotek Cytation3 imaging reader. To perform the Brdu assay, cells were seeded on coverslips in a 24-well plate and pulsed with 6 µg/ml BrdU (B5002, Sigma, USA). After 2 h, the cells were fixed and stained with anti-BrdU antibody (Millipore, Burlington, MA, USA) as described. To perform the colony formation assays, transfected cells were seeded in 6-well plate at a density of 100–

250 cells per well and incubated for 2 weeks. The cells were fixed and stained in a dye solution containing 0.1 % crystal violet and 20 % methanol. The number of colonies with more than 50 cells was counted.

**Wound healing, cell migration and invasion assay**

The effects on migratory function of miR-107 overexpression and KIF23 knockdown were determined by evaluating cellular migration after scratching of a confluent monolayer of cells. Cells were seeded into 6-well plates and transfected with either siRNA, miRNA mimic, miRNA antagomir or control. After 48 h, the cells were starved for additional 24 h before the pre-treatment of the cells with 10 µg/ml Mitomycin C (M0503, Sigma Aldrich, St. Louis, MI, USA) for 3 h. Mitomycin C is a cell cycle antagonist being used to measure migration without the influence of cell proliferation. The cells were washed three times followed by the scratch drawn using a 200-µl pipette. Migration was photographed at the indicated time points after scratching using an inverted microscope (Leica, Wetzlar, Germany) and the percent wound closure was calculated for five randomly chosen fields.

Migration assay was also confirmed by performing by using Transwell insert chambers (24-well plate, 8.0 µm pore size, Corning, Corning, NY, USA). $1*10^5$ transfected cells suspended in serum-free medium were placed in the upper chamber 48 h after transfection while the lower chamber was filled with 700 µl of DMEM with 10 % FBS as chemo-attractant. After incubation for 24 h at 37 °C, the non-migrating cells in the upper chamber were removed by a cotton swab and lower surface of the chamber was fixed and stained with 0.1 % crystal violet. For invasion assay, the cells were seeded in insert chambers pre-coated with a layer of diluted basement membrane matrix Matrigel (Corning, Corning, NY, USA) and the same procedure was followed as above. Migrating or invading cells were scored by counting at least 5 fields per membrane under a light microscope.

**Cell culture and transfections of miRNA inhibitors and siRNA.**

Liver cancer cell lines Hepa1–6, Huh-7, Hep3B and HepG2 were obtained from the American type culture collection (Manassas, VA, USA). Hepa1c1c7 cell line was a kind gift from Dr. Irina Lehmann (Helmholtz Centre for Environmental Research GmbH-UFZ, Leipzig Germany). All cells were cultured in DMEM medium (Pan Biotech, Germany) containing 10 % fetal bovine serum (Invitrogen, CA, USA) and 100 unit/ml of penicillin/ streptomycin (Invitrogen, CA, USA). Miscript miRNA mimics, siRNAs, miRNA antagomiRs and negative control were purchased from Qiagen (Hilden, Germany). siRNA or miRNA mimics and All Stars negative control (Control siRNA) were used at final concentrations of 50 nM. miR-107 inhibitor and miScript Inhibitor Negative control were used at final concentrations of 100nM. Cells were transfected using RNAiMax (13778075; Invitrogen, CA, USA) according to manufacturer's instructions and let recover for 72 h before analysis. Transfection of pEGFP-hsa-107 and pEGFP-CTRL in Huh-7 were transfected with linear 25 kDa polyethyleneimine (PEI, Polysciences Inc., Warrington, PA, USA) by using a 3:1 ratio of PEI/DNA as previously described [1]. The complete list of miRNA mimics/inhibitors and siRNAs used in this study is available in the **Supplementary Table 9**.

**Bioinformatic analysis of Kif23/miR-107 interaction and dual luciferase assay**

RNA22 [6] was used to predict potential miR-107-binding sites in the 3′UTRs of human and mouse Kif23. Full-length 3′UTRs were amplified using an Expand High Fidelity PCR System (Roche, Mannheim, Germany, 11732641001) as indicated by the vendor. Primers used for 3′UTR amplification are listed in **Supplementary Table 8**. The resulting 3′UTRs were cloned into pMIR(+) and pMIR(-) reporter vectors as previously described [7]. Constructs carrying the 3′UTRs were transiently co-transfected with vectors carrying Renilla luciferase and either miR-107 mimic or scramble oligos (SCR) into HEK293 cells with Lipofectamine RNAiMAX (Thermo Fisher Scientific, Meerbusch, Germany, 13778030) according to the manufacturer's instructions. A dual-luciferase reporter assay (Promega, Walldorf, Germany, E1910) was conducted in accordance with the manufacturer's instructions. The chemiluminescence assay was performed in white opaque 96-well plates with 50 µL of cell lysate and 50 µL of both LARII reagent (Firefly luciferase substrate) and Stop and Glo reagent (Renilla luciferase substrate), in a GloMax Multi Plus Multiplate Reader (Promega, Walldorf, Germany) according to the preset protocol with an integrity time of 10 s for each read. Data were normalized by calculating ratios of firefly/Renilla activities to correct for possible variations in transfection efficiencies.

**HDTVi tumor models and genetic mouse models for liver cancer**

shRNA sequences were generated using the SplashRNA online tool [8] and cloned via EcoRI and XhoI into a doxycyclin inducible retroviral expression vector (RT3GEPIR) including a mirE based miRNA cassette as previously described [9]. To test the efficacy of the distinct shRNA sequences, Hepa1-6 cells transfected with the RT3GEPIR vectors carrying the shRNA/pre-miR-107 and Kif23 expression was assessed at RNA and protein levels 72 hours after transfection.

The most efficient shRNA sequences, the pre-miR-107 sequence and the mirE cassette were excised from the RT3GEPIR vectors via XhoI and MluI and subcloned in the CamiN-mA-E transposon vector, encoding for Myc/NRas oncogenes, cut with XhoI and AscI. To generate the "positive control" CamiN-miRE vector the sequence for the mir30e-stem loop (mirE) from the RT3GEPIR vector was directly subcloned into the CamiN-mA-E vector as indicated above. To perform hydrodynamic tail vein injection (HDTVi), 7 week old male C57BL/6j mice were anesthetized with Isofluran, and a solution composed of - 25 µg CamiN-mA-E transposon vectors, 5 µg SB13 transposase in 0.9%NaCl - 10% v/w body weight - was injected into the tail vein within 5 seconds (as described [10]). Mice were housed under pathogen free conditions and fed with normal diet and were euthanized 7 weeks after HDTVi.

All animal experiments were approved and conducted according to the local authority (Regierungspräsidium Dusseldorf, NRW, Germany). This method was performed to test the knockdown efficiency of shRNAs by isolating total RNA using Qiazol (Qiagen, Hilden, Germany) followed by chloroform purification and purification via RNeasy columns (Qiagen, Hilden, Germany). cDNA synthesis and for mRNAs and miRNAs were performed as previously described [2, 7] and qPCR was conducted with SYBR Green I (GO-Taq PCR Master mix, Cat: A6002; Promega, Walldorf, Germany) in a ViiA7 real-time PCR machine (Applied Biosystems). Relative mRNA expression was calculated with the ΔΔCT method [5]. The list of primers and shRNA sequences used for this experiments are listed in **Supplementary Table 8**. The three different mouse models for liver cancer, the generation and analysis of these animals [i.e., liver tumors driven by diethylnitrosamine (DEN), lymphotoxin alpha and lymphotoxin beta (AlbLTα/β) and Myc-driven (Tet-O-Myc)] were published previously [9].

**Serum preparation and analysis**

Sera were prepared from mouse blood by using micro sample tube clotting activator gel (Sarstedt, Nümbrecht, Germany, 41.1500.005) according to manufacturer's instructions. The levels of aspartate aminotransferase (AST), alanine aminotransferase (ALT) and glutamate dehydrogenase (GLDH) the serum of mice were measured by standard procedures in our laboratory by using a Pentra C400 Clinical Chemistry Analyzer (Horiba Medical, Kyoto, Japan).

**Preparation, Staining and ISH analysis of FFPE slices**

Liver tissues were fixed in Paraformaldehyde (4%) and paraffin embedded. FFPE blocks were sliced (2 µm sections) and stained with either hematoxylin and eosin (H/E) to visualize liver structure or Sirius red to visualize collagen deposition. Image acquisition was with an Aperio AT2 slide scanner (Leica, Wetzlar, Germany). Digital images were visualized and analyzed with the Aperio ImageScope software (version 12.4.6; Leica, Wetzlar, Germany).

**Immunofluorescence, IHC staining and analyses**

Cells were grown on coverslips and transfected as indicated above. At the end of treatment cells were fixed in 4% PFA, blocked with normal goat serum diluted in 0.1 % Triton X-100. The cells were washed three times with PBS and incubated overnight at 4 °C with either of the following primary antibodies: Ki67 (NCL-Ki67p, Leica Microsystems), α-Tubulin (T6074, Sigma Aldrich, St. Louis, MI, USA), or KIF23 (120241–1-AP, Proteintech, Rosemont, IL, USA) and treated with secondary antibodies anti-rabbit A555 (A31572, Invitrogen, CA, USA), and anti-mouse A488 (A21202, Invitrogen, CA, USA). Image acquisition was performed at a magnification of 20 x with a Zeiss Axio Imager.Z1 microscope, Axiocam MRm and HRc cameras using Axiovision 4.8 software (Carl Zeiss, Oberkochen, Germany). For Immunohistochemistry (IHC), Paraformaldehyde (4%) fixed and paraffin embedded liver tissue sections (2µm) were stained in an HRP-based DAB staining (DAB Quanto, Thermo Fisher Scientific, Meerbusch, Germany) with various primary and secondary antibodies. The following antibodies were used: Anti F4/90 (Abcam, 1:300), Anti-Rat IgG (Vector), Anti-CD3 (Thermo Fisher Scientific, Meerbusch, Germany. 1:500), Anto-Rabbit IgG (Thermo Fisher Scientific, Meerbusch, Germany). Image acquisition was performed with an Aperio AT2 slide scanner (Leica, Wetzlar, Germany), acquired images were visualized using ImageScope (Leica, Wetzlar, Germany, Version 12.4) and analyzed using QuPath (Version 0.4.3).

For Immunofluorescences (IF), cells were grown on cover slips and fixed with 4% PFA. Following fixation and permeabilization, proteins of interest were stained by incubation with primary and secondary antibodies, while DNA was stained using DAPI. Stained cover slips were mounted and images were captured using a CZ Axio Observer Z1 microscope (Carl Zeiss, Oberkochen, Germany) equipped with a 63x objective and acquired with ZEN blue light (version 3.6).

**Western Blot**

Proteins were isolated using RIPA buffer with protease inhibitor cocktail (Complete Protease Inhibitor Cocktail, Roche, Mannheim, Germany, 04693116001), and concentrations were measured using the Qubit Protein Assay Kit (Thermo Fisher Scientific, Meerbusch, Germany, Q33211) according to the manufacturer's instructions. Western blot analyses were carried out using 20 µg of total proteins. Protein lysates were loaded together with 10 µL of Protein Ladder (BioRad, Dusseldorf, Germany, 161-0373) on 10% or 12% SDS polyacrylamide gels and transferred to nitrocellulose membranes using semidry blotting systems according to standard protocols. Membranes were blocked with 5% milk powder (Carl Roth, Karlsruhe, Germany, T145.3) in Tris-buffered saline with Tween20 (TBST) for 1 h at RT or overnight at 4 °C, followed by incubation with a horseradish peroxidase (HRP)-coupled antibody against rat albumin (Bethyl Laboratories, Montgomery, TX, USA A110-134P) for 2 h at room temperature. Chemiluminescence was detected with ECL Western blotting Substrate (Promega, Walldorf, Germany, W1001) using the ChemiDoc MP Imaging System (BioRad, Dusseldorf, Germany). Signal intensities of

Western blot protein bands were analyzed using Image Lab (BioRad, version 6.0.1). Antibodies used in analysis by Western blot: Anti-KIF23 (1:100, Abcam Cambridge, MA, USA, ab174304), Anti-KIF23 (1:100 dilution, Proteintech, Rosemont, IL, USA, 120241–1-AP), Anti-KIF23 (1:100 dilution, Santa Cruz, Dallas, TX, USA, sc-390113), anti-GAPDH (1:5000, ABD Serotec, Raleigh, NC, USA, MCA 4739), anti-Albumin antibody (1:2000 dilution, Abcam, Cambridge, UK, ab207327), anti-bActin (1:2500, Cell signaling, Danvers, MA, USA, 4967S).

#### **Supplementary References**References
